# Single-cell RNA sequencing clarifies dermal fibroblast subset representation *in vitro* and reveals variable persistence of keloid disease-associated features

**DOI:** 10.1101/2025.08.11.668109

**Authors:** Amy Lock, Elena M Drudi, Dasha Freydina, Brian M Stramer, Franziska Denk, Tanya J Shaw

## Abstract

*In vitro* models of scarring and fibrosis are essential to improve our understanding of disease mechanisms and ultimately develop much-needed effective therapeutic strategies. This is particularly true for keloids, the example of pathological scarring exploited in this study, as there is no animal model. Our emerging appreciation of fibroblast heterogeneity from single cell RNA sequencing (scRNA-seq) information leaves a knowledge gap about what is represented in typical fibroblast cultures. Specifically, it is important to know whether quantitative differences in fibroblast subtypes observed in pathological tissues are represented and/or whether disease-associated molecular alterations of subtypes are maintained.

This study performed scRNA-seq on patient-matched keloid and normal adjacent dermis immediately following surgical removal, which was compared to sc- and bulk-RNA-seq on primary dermal fibroblast cultures from the same samples after 4+ passages. Freshly dissociated tissue showed anticipated differences in cell proportions in keloid versus normal skin; however, comparably for both tissue types, there was an assimilation of fibroblast subtypes after culture. Cultured cells clustered conspicuously from the original populations, with evidence of only minor heterogeneity persisting. Cells displayed, to varying degrees, elements of each of the original subset signatures, with FAP^+^/SFRP2^+^ mesenchymal features the strongest. Pseudo-bulk analysis of mesenchymal subpopulations *ex vivo* showed cell-intrinsic keloid versus normal skin transcriptional differences consistent with current disease understanding; however, only a subset of these persisted *in vitro*. Cell-cell communication analysis provides potential strategies to maintain specific cell populations and their *in vivo* phenotypes. As an example, we report that culture with ascorbic acid (stimulating cell-derived extracellular matrix) enriched the mesenchymal signature. The data presented herein provide resources supporting greater understanding of, and strategies to refine, essential human fibroblast culture models.

## INTRODUCTION

Keloids are a type of pathological scar characterized by abnormal collagen deposition and expansion beyond the wound boundary. They continue to grow over time, recur after resection and do not regress spontaneously (Chike-Obi et al., 2009). As such, these scars can result in sizeable masses with significant aesthetic concerns (Kim et al., 2022). Keloids are highly fibrotic and share many characteristics of fibrosis found in other areas of the body; such characteristics include a highly aligned architecture (Kenny et al., 2023), fibroblast accumulation and excessive and high collagen I:III ratio (Friedman et al., 1993), leading to stiff tissue that is no longer able to function as adequately as its normal counterparts. Together with the ease of access to these surface-based lesions, keloids are an excellent model condition to study the biology of fibrosis.

Recently, several single-cell RNA sequencing (scRNA-seq) studies have compared cell type variety and abundance in keloids compared to normal scar and skin (Deng et al., 2021, Direder et al., 2022a, Liu et al., 2021). These studies, individually and when considered collectively (Direder et al., 2022b), have revealed distinct cell subsets to be enriched in keloids, such as extracellular matrix (ECM)-producing fibroblasts, amongst others. Building on these findings, developing *in vitro* models to study these cell types and their mechanisms is essential for advancing keloid research. As there is no suitable animal model that faithfully recapitulates keloid disease, the ability to expand primary cells is paramount; but there is concern about loss of *in vivo* phenotype (Neumann et al., 2010, van Vijven et al., 2021, Yang et al., 2018) and phenotypic drift (Machaliński et al., 2020). The recent scRNA-seq publications illustrating diverse cell composition in keloids indicate that it would be beneficial to be able to isolate and maintain particular (sub)populations for detailed mechanistic work.

In this study, we investigated the transcriptional changes that primary dermal cells isolated from keloid or adjacent skin undergo during standard fibroblast culture conditions by scRNA-seq. We show that passage (P)4 cells are transcriptionally different to their *in vivo* (P0) derivatives, with changes in cell subset identity and proportions, and show variable effects on disease-associated transcriptional differences between keloid and normal fibroblasts. We also provide insight into how culture conditions can be altered based on cell-cell communication analysis to shift cultured fibroblasts towards a particular (mesenchymal-like) phenotype.

## RESULTS

### Dermal cell subsets are transcriptionally different by the 4th passage

To improve our understanding of the extent of changes to dermal cell subsets over passage, we performed scRNA-seq to compare acutely isolated cells (P0) with those cultured over four passages (P4). Dermal cells were extracted from an abdominal keloid scar and patient-matched adjacent normal skin. Consistent with published studies (Deng et al., 2021, Direder et al., 2022a, Liu et al., 2021), unsupervised clustering revealed nine cell types (Figure 1a, c). In this particular sample, smooth muscle cells (SMC)/myofibroblasts represented the majority of keloid cells, with fibroblasts and endothelial cells representing similar proportions to each other (∼10%). Surprisingly, the most numerous cell types in normal adjacent skin were T-cells and NK cells (>50%; far exceeding that expected of normal skin), potentially reflecting an active immune response at the lesion margin in this patient.

**Figure 1.**
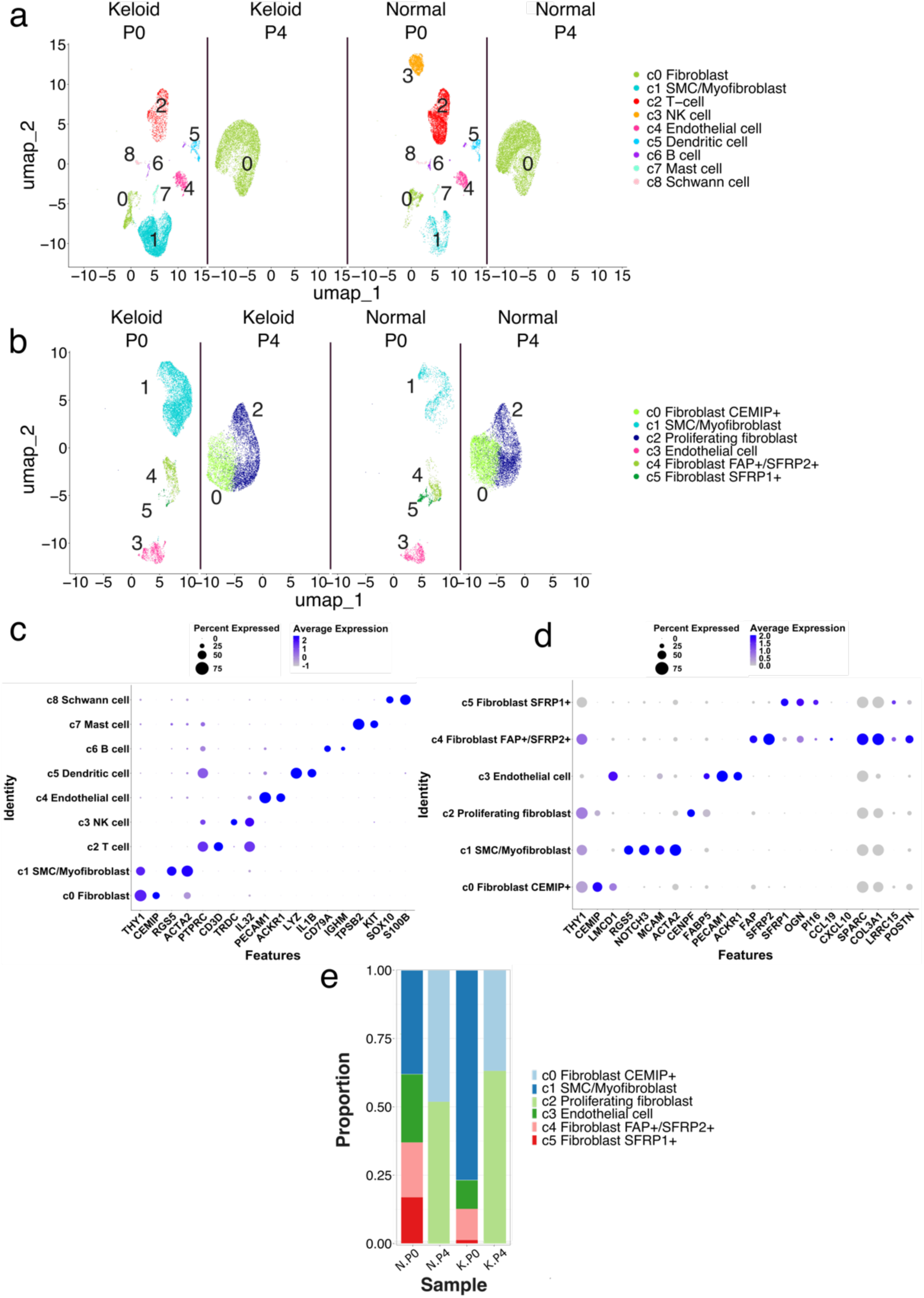
Single-cell RNA sequencing reveals dermal cell subsets are transcriptionally different by the 4^th^ passage in culture. (a-b) Sample-separated UMAPs of the unsupervised clustering of (a) all cells (N=34,948) revealed 9 clusters and (b) stromal cells only (N=25,956) revealed 6 clusters. P0 = passage 0, P4 = passage 4. (c-d) Expression of cluster marker genes across each cluster. Blue color gradient represents average expression of given gene. Circle size represents the percentage of cells within a cluster expressing a given gene. (e) Proportion of each stromal subset out of total stromal cells for normal primary dermal cells at passage 0 (N.P0) and passage 4 (N.P4) and keloid dermal cells at passage 0 (K.P0) and passage 4 (K.P4).

There were stark differences in the cell types present at P4 compared to P0 for both keloid and normal skin, with only cells identified as fibroblasts still present (Figure 1a). Selective unsupervised re-clustering of the stromal cells across time-points revealed six sub-clusters, which were annotated as: CEMIP^+^ fibroblast (c0), SMC/myofibroblast (c1), proliferating fibroblast (c2), endothelial cell (c3), FAP^+^/SFRP2^+^ fibroblast (c4), and SFRP1^+^ fibroblast (c5) (Figure 1b, d). Both normal and keloid P4 cells were remarkably transcriptionally different to the four cell subsets present at P0, with only two distinct clusters observed: CEMIP^+^ fibroblasts and proliferating fibroblasts (Figure 1e). These cell populations were additionally annotated according to “atlas datasets” describing universal fibroblasts in health and disease (Buechler et al., 2021, Korsunsky et al., 2022). Using the markers of fibroblasts in healthy tissues, the FAP+/SFRP2+ subset was delineated by SPARC/COL3A1 whereas no subsets were marked by CCL19/CXCL10, indicating a lack of a proinflammatory subset in this particular sample (Figure 1d). Interestingly, the FAP+/SFRP2+ subset expressed key markers also observed in universal perturbed fibroblasts (LRRC15, POSTN) (Figure 1d).

### Passaged cells represent transcriptional assimilation of *in vivo* subsets

We next increased the granularity of our analysis to investigate the potential for subset heterogeneity after culture; selective re-clustering of P4 cells revealed four subsets (Figure 2a), annotated as CEMIP^+^, proliferating, KRT19^+^ and SRGN^+^ (Figure 2b). Subset proportions varied slightly between groups, with cultured keloid cells containing more proliferating, KRT19^+^ and SRGN^+^ cells (Figure 2c). To determine which P0 cell clusters were best represented by the P4 cultures, if any, cluster scores were calculated based on ‘signatures’ (Supplementary Data 1) using the UCell package (Figure 3). Interestingly, CEMIP^+^ fibroblasts shared a degree of transcriptomic similarity across all P0 fibroblast and mural (SMC) subsets, indicating they may represent a common intermediate, or a culture-induced mix (Figure 3a, e-g). Proliferating and KRT19^+^ fibroblasts appeared to be culture-specific phenomena, sharing very little similarity to any P0 subset (Figure 3b-c). P4 SRGN^+^ cells, though only a small population, shared the most similarity to P0 endothelial cells, a cell type which classically requires specialized culture supplements to be maintained (Figure 3d). Together, these data suggest that while some fibroblast states may persist or converge during culture, others - particularly the proliferating, KRT19^+^, and SRGN^+^ subsets - likely emerge as culture-induced adaptations.

**Figure 2.**
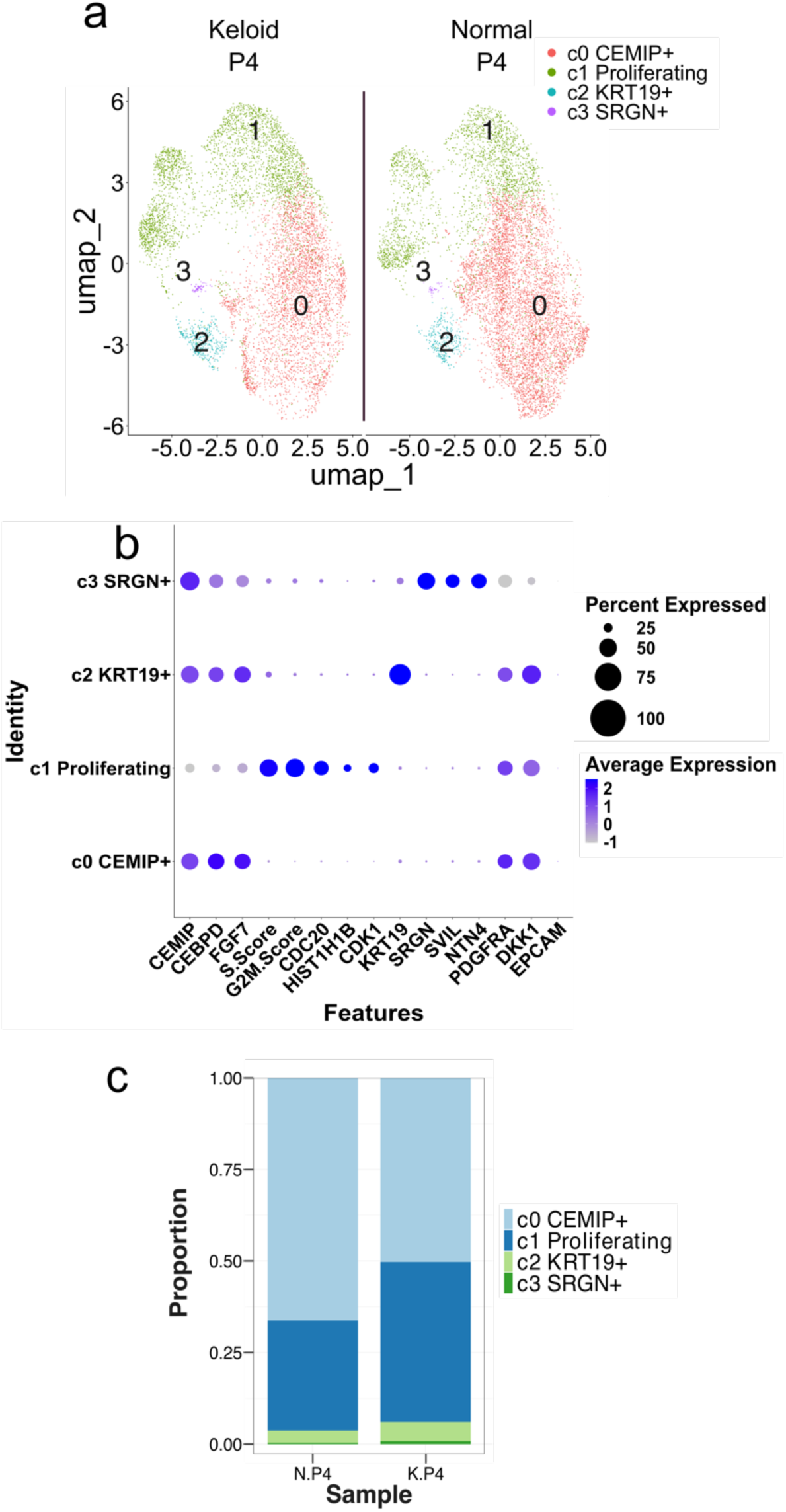
Passage 4-only sub-clustering reveals stromal heterogeneity still present at later passage. (a) Sample-separated UMAPs of the unsupervised clustering of 15,255 cells revealed 4 clusters. (b) Expression of cluster marker genes across each cluster confirming cell identity. Blue color gradient represents average expression of given gene. Circle size represents the percentage of cells within a cluster expressing a given gene. (c) Proportion of each subset out of total cells for normal (N.P4) and keloid primary dermal cells (K.P4) at passage 4.

**Figure 3.**
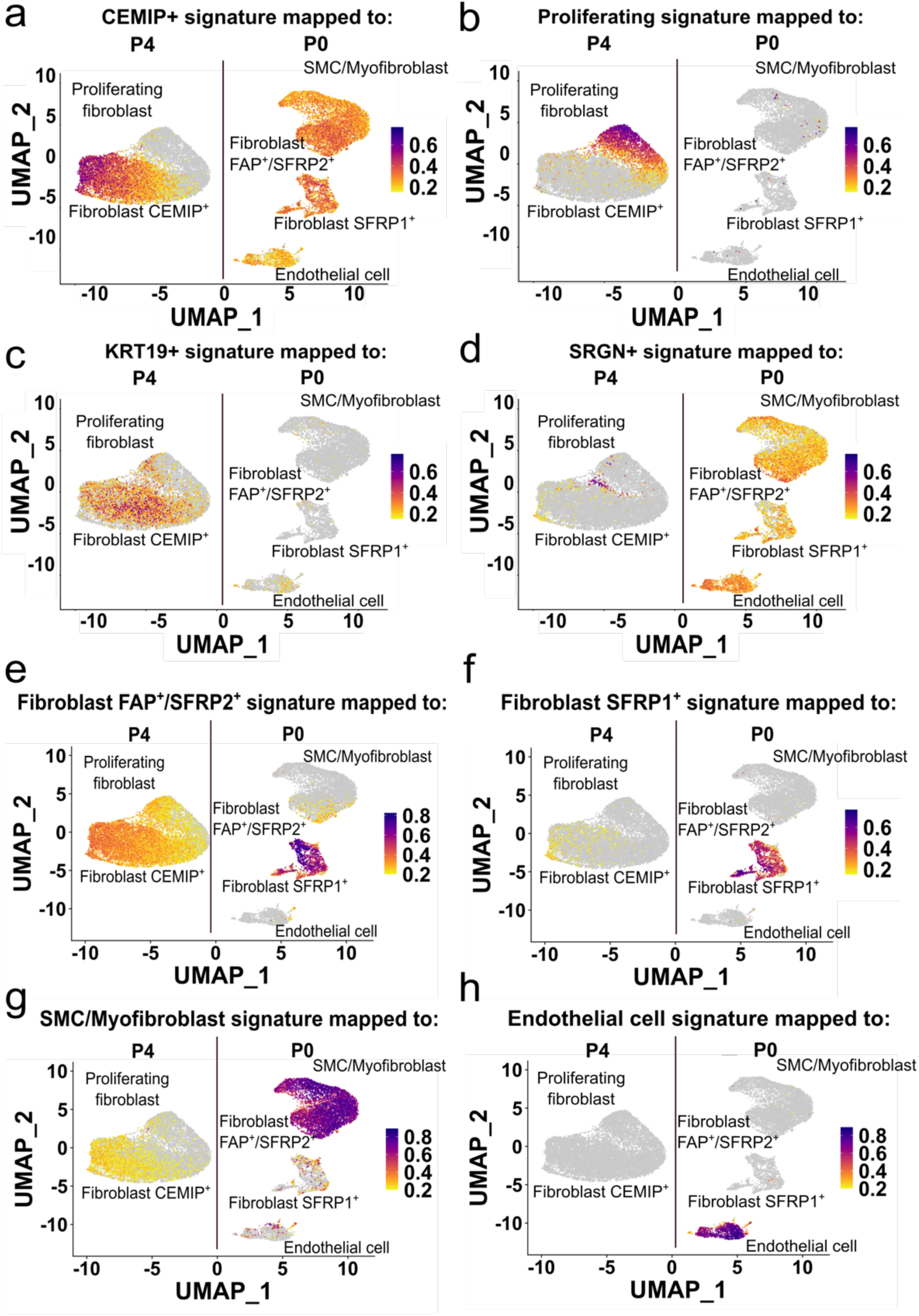
Passage 4 cells represent an amalgamated or common intermediate phenotype of passage 0 stromal cell subsets. (a-h) Feature plots showing the P4 cluster scores of (a) CEMIP+, (b) proliferating, (c) KRT19+, (d) SRGN+, and P0 cluster scores of (e) Fibroblast FAP+/SFRP2+, (f) Fibroblast SFRP1+, (g) SMC/Myofibroblast and (h) Endothelial cell subsets mapped to the UMAP of the P0/P4 stromal dataset. Grey-purple color gradient represents increasing cluster score (gene signature similarity) of the given subset, per each cell. Plots created using the scCustomize package.

As this sequencing study was based on single technical repeats, the observed changes in cell composition after culture were further investigated by analysing bulk-RNA-seq for expression of subset transcriptional marker genes in four further higher-passage samples (P8-9) from the same individual. In keeping with our scRNA-seq results, P0 cluster markers were largely decreased or lost in >P4 samples, across both keloid and normal groups (Figure 4a). In keeping with our scRNA-seq data, FAP^+^/SFRP2^+^ subset markers were the most maintained, highlighting this subset as the most represented at later passage. Furthermore, to align our data to widely used fibroblast nomenclature, we ascertained subset marker expression first introduced by Solé-Boldo et al (2020) (Figure 4b). As expected, some markers were decreased or absent in P8/P9 samples, with secretory-papillary and secretory-reticular subsets most widely represented. To validate our findings in additional primary patient samples and cultures, we investigated passage changes in cell composition using transcript abundance of marker genes as a surrogate assessed by qPCR in four unmatched samples of normal and keloid skin, again comparing P0 to P4 (Figure 4c-f). As described above, many P0 subset markers were decreased in P4 samples, and this was observed in both normal and keloid dermal cell cultures. However, there was interesting variability; for example, SFRP2 expression was downregulated, whereas FAP expression was maintained contrary to the P0/P4 scRNA-seq (Figure 4d), indicating that transcriptional profile changes are not global and/or may be subject to donor variability. Indeed, qPCR for some P4 cluster markers validated our sequencing data (CEMIP, CDK1, KRTAP1-5), while others were absent (CEBPD, MKI67) or showed unexpected expression patterns (SRGN) (Supplementary Figure 1).

**Figure 4.**
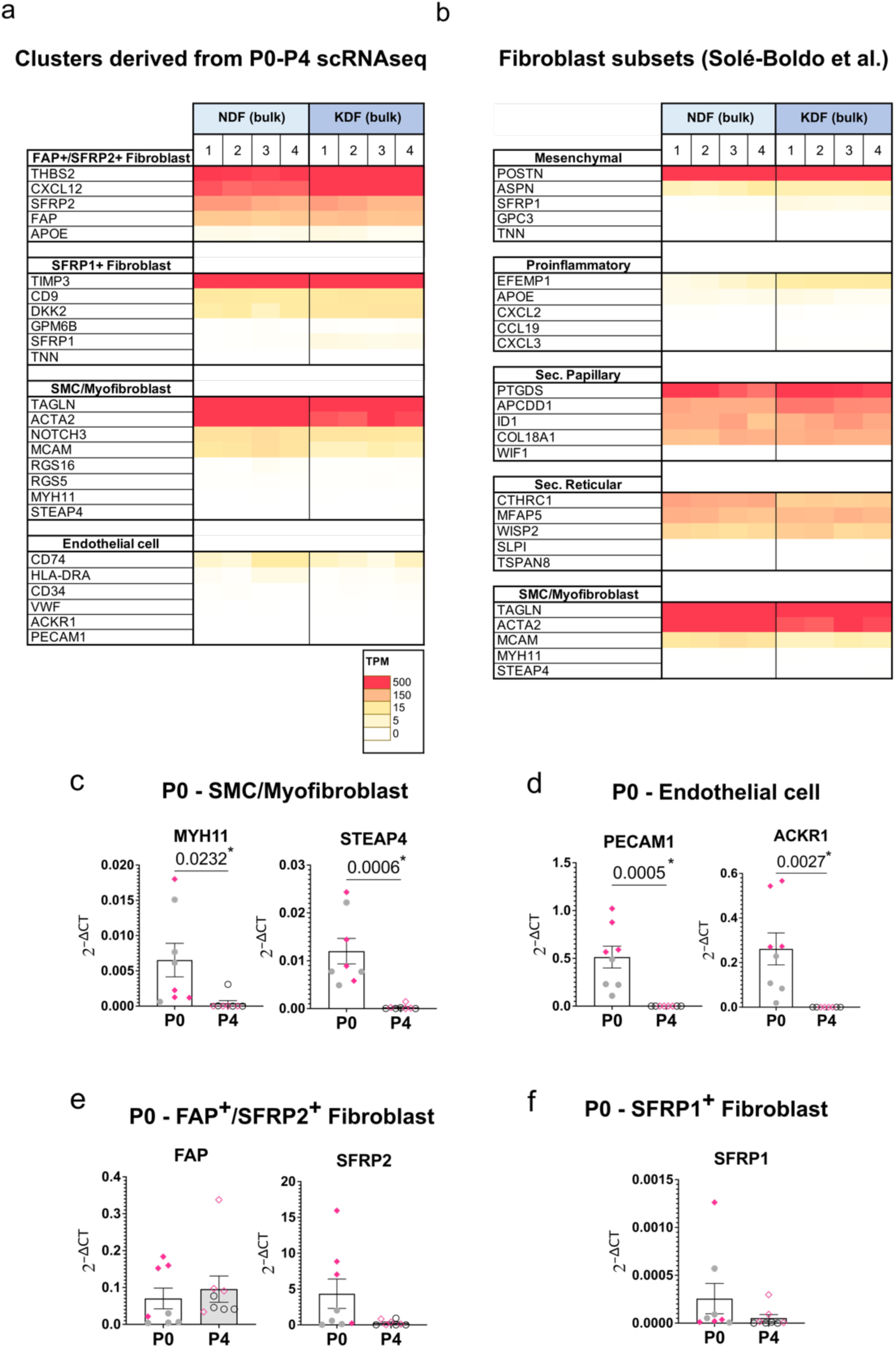
Confirmation of assimilated phenotype in further expanded dermal cell samples. (a, b) Raw transcripts per million (TPM) expression of (a) P0 stromal or (b) skin fibroblast subset markers (Solé-Boldo et al., 2020) in bulk RNA sequenced normal and keloid fibroblasts from P8-9. (c-f) Expression of passage (P) 0 stromal subset markers in P0 (white bar) and P4 (grey bar) normal (grey circles) and keloid (pink diamonds) dermal cells. N=8/group, with each dot derived from a different donor. qPCR expression normalized to reference gene, GAPDH. Bars represent mean +/-SEM. Statistical test: Unpaired T-test; p-values are displayed on the graph.

### Differential expression in keloid mesenchymal fibroblasts is not represented in passaged cells

Keloids represent an extreme fibrotic condition and the fibroblast subsets that they contain differ not only in proportion, but also in their transcriptional profiles compared to normal skin. Integration of keloid scar and normal skin samples from three published scRNA-seq datatsets (Deng et al., 2021, Direder et al., 2022a, Liu et al., 2021) showed that mesenchymal fibroblasts are significantly increased in keloids compared to other subsets (Figure 5a). Focusing on this subset, we used pseudo-bulk counts and carried out differential gene expression analysis to identify the keloid-specific phenotype (Figure 5b). In total, 227 genes (log_2_FC<-0.5 or >0.5, padj<0.05) were significantly differentially regulated in mesenchymal fibroblasts from keloids (n = 11 samples) compared to normal skin (n = 4 samples), including known cartilage-related genes such as ASPN and BGN, as previously noted in proteomics analysis (Barallobre-Barreiro et al., 2019), and other markers (e.g., MDK, SULF2) previously shown to be specific to keloid mesenchymal fibroblasts compared to normal scar (Deng et al., 2021) (Supplementary Data 1). To assess whether keloid-specific differences persist through passaging, we sought to identify how many of the genes regulated in the *ex vivo* mesenchymal fibroblast data can still be observed in bulk RNA-sequencing of passage 8 and 9 fibroblasts (Figure 5c). Of the 227 differentially expressed *ex vivo* genes, 211 remained expressed at P8/9, but only 26 and 8 remained differentially up- or down-regulated (adj. p < 0.05), suggesting the loss of keloid mesenchymal fibroblast phenotype over passage or, alternatively, acquisition of fibrotic features by normal cells. Of note, the bulk-RNA sequencing still uncovered significant differential expression between keloid and normal fibroblasts (296 DEGs; log_2_FC<-0.5 or >0.5, padj<0.05), indicating disease-related differences continue or are highlighted through culture (Supplementary Data 1).

**Figure 5.**
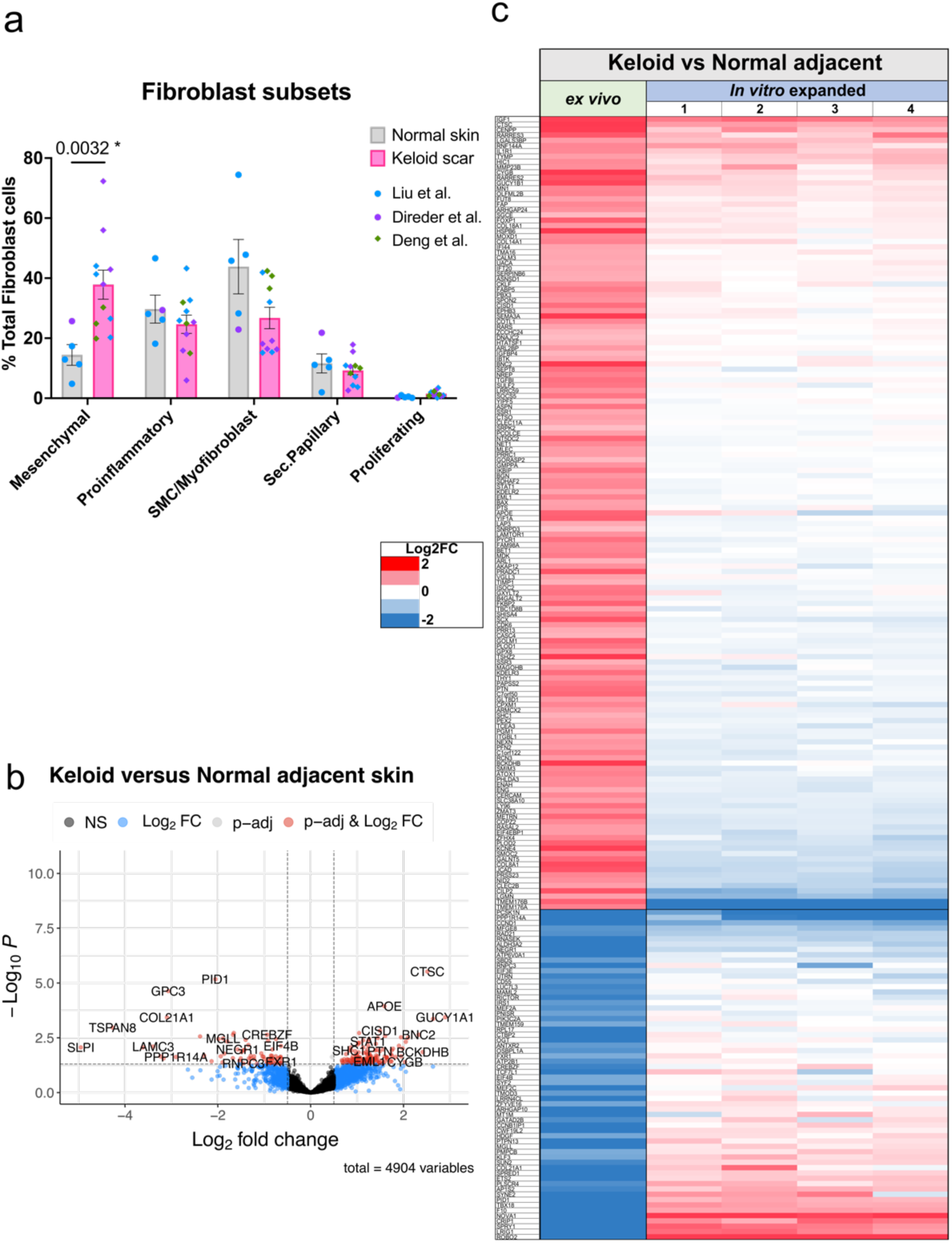
Disease-associated differential gene expression in keloid versus normal mesenchymal fibroblasts is diminished in passaged cells. (a) Proportion of fibroblast subsets out of total fibroblast cells for normal skin (grey, N=5 biopsies) and keloid scar (pink, N=11 biopsies from different individuals). Data derived from Liu et al. (blue points), Deng et al. (green points), and Direder et al. (purple points) scRNA-seq datasets. Statistical test: unpaired, non-parametric T-test (Mann-Whitney) for which significant p-values are displayed on the graph. (b) Volcano plot displaying differentially expressed genes for keloid (N=11 biopsies) versus normal adjacent skin (N=4 biopsies) mesenchymal fibroblasts. Adjusted P cutoff: <0.05, Log2FC cutoff: >0.5 or <-0.5. Plots created using Enhanced Volcano R package. (c) Heatmap showing Log2 fold change (Log2FC) values of differentially expressed keloid mesenchymal fibroblasts genes in *ex vivo* scRNA-seq keloid mesenchymal fibroblasts and bulk RNA sequenced keloid fibroblasts from P8-9 (1-2 = P8, 3-4 = P9).

### Strategies for preservation of fibroblast cell subsets across passage

To overcome the lack of subset longevity in culture, we explored the possibility of triggering certain signaling pathways to sustain cell phenotype. We reasoned that heavily utilized incoming signaling pathways *in vivo* (i.e. at P0) may be important; therefore, we used the cell-cell communication package CellChat DB (Jin et al., 2021) to identify top incoming (and outgoing) signaling pathways (Figure 6a-b). Mesenchymal fibroblasts showed several enriched pathways relating to matrix interaction (collagen, FN1), tissue regeneration (PTN, MDK) and mesenchymal lineage (periostin, CD99). The strength of matrix interaction and the autocrine signaling loop of periostin (POSTN) via the target receptor pair ITGAV and ITGB3 (Figure 6a-b), hinted that interaction with the ECM may be responsible for driving the mesenchymal phenotype, and mimicking this trigger in culture may support their maintenance. We evaluated whether stimulating the cells to lay down their own ECM during culture would shift their phenotype. We bulk-RNA sequenced P8/P9 keloid and matched-normal skin cells that were cultured for 5 days using standard fibroblast conditions with or without supplementation with ascorbic acid, a necessary co-factor for collagen synthesis and other ECM component deposition (Kao et al., 1990, Phillips et al., 1994) (Figure 6c). Using the gene signatures of fibroblast subsets from our integrated scRNA-seq analysis (Supplementary Data 1), we showed the mesenchymal signature was stronger in samples cultured with ascorbic acid (Figure 6d-h), indicating ECM production and interaction shifts fibroblasts towards a more mesenchymal-like phenotype.

**Figure 6.**
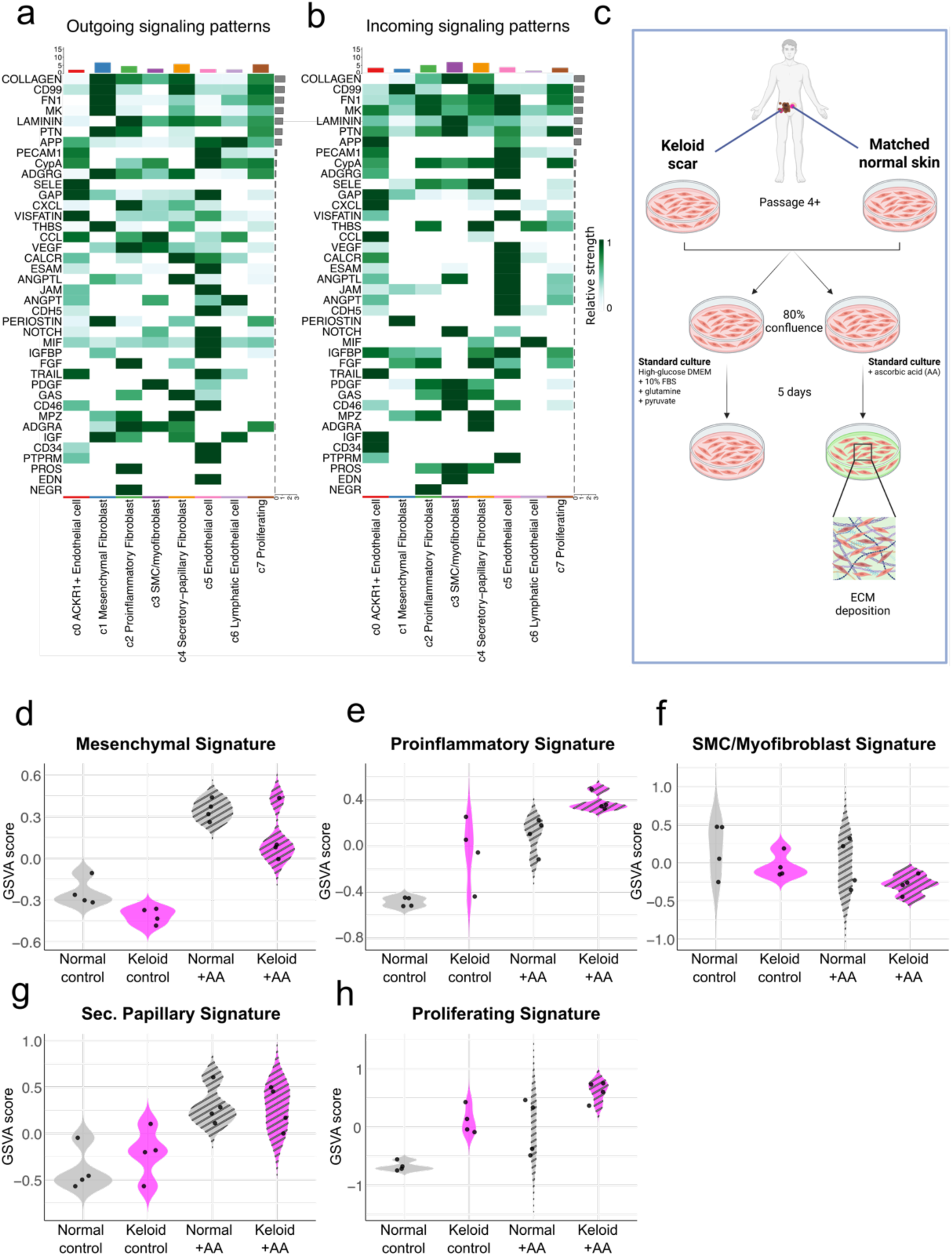
Addition of ascorbic acid to fibroblast culture is sufficient to shift fibroblasts towards a more mesenchymal phenotype. (a-b) Heatmap showing the top 25 (a) outgoing and (b) incoming signaling pathways of stromal cell subsets in keloid scars. The green color bar represents the relative signaling strength of pathways across cell subsets, with darker green indicating higher strength (row-scaled values). The colored bar plot at the top shows the total signaling strength of each cell subset, summarizing all pathways. The right grey bar plot indicates the total signaling strength of each pathway, summarizing all cell subsets. Generated using the CellChat package in R. (c) Schematic outlining experimental procedure. Matched keloid and normal skin cells at P8-9 were seeded and grown in standard fibroblast culture conditions with or without ascorbic acid for 5 days. (d-h) Violin plots showing gene set variation analysis (GSVA) scores of (d) mesenchymal, (e) proinflammatory, (f) smooth muscle cell (SMC)/myofibroblast, (g) secretory (Sec.) papillary and (h) proliferating fibroblast gene signatures in bulk RNA sequenced normal and keloid fibroblasts with (+AA) or without ascorbic acid (control). GSVA completed using GSVA package in R.

## DISCUSSION

Here we aimed to address key unknowns about primary dermal fibroblast culture, specifically the cell subtypes represented, cell heterogeneity (if any), the transcriptional changes induced by culture, and whether disease-associated molecular alterations are maintained.

As our understanding of fibroblast subtypes based on scRNA-seq across health and disease is still relatively new (Buechler et al., 2021, Korsunsky et al., 2022), culture-induced changes at this level have only been minimally considered. We used scRNA-seq to examine the changes in cell representation and proportion using a patient-matched pair of normal and keloid dermal cells immediately following tissue dissociation (at P0) and after four passages in standard culture conditions (P4). Due to the within-subject design, we could isolate changes due to the culturing process alone. As expected, we found non-fibroblast cells were not retained, as we specifically enriched for plastic-adherent cells in conditions typical for fibroblast culture (high-glucose DMEM with 10% serum; no specialized growth factors or substrates). Next, focusing on stromal cells only, we observed stark changes; at P0, three distinct fibroblast subtypes were present: FAP^+^/SFRP2^+^, SFRP1^+^ and SMC/myofibroblasts, with FAP^+^/SFRP2^+^ akin to the SPARC^+^/COL3A1^+^ universal fibroblast subtype, while SFRP1^+^ correspond to the PI16^+^ universal subset (Buechler et al., 2021, Korsunsky et al., 2022). Both FAP^+^/SFRP2^+^ and SFRP1^+^ subsets also expressed markers of perturbed/pathological fibroblasts (Buechler et al., 2021): LRRC15/POSTN/COL3A1 and LRRC15/PI16, respectively (Supplementary Figure 2). Yet by P4, cell phenotypes had dramatically changed and no longer clustered with P0 counterparts. P4-only clustering revealed four distinct clusters: CEMIP^+^, proliferating, KRT19^+^ and SRGN^+^, and this was equal for both normal- and keloid-derived cells. Although CEMIP^+^ and proliferating cells constitute the overwhelming majority (>90%), this diversity shows the cultures are not completely homogeneous as has been previously suggested (Dirand et al., 2023, Harper, 1989, Ladin et al., 1998). Moreover, their functional differences and plasticity remain to be determined. Mapping the P4 cluster signatures to P0 subsets to understand which of the original populations are represented in the culture showed their markers dispersed across clusters, suggesting they represent a convergence, or common intermediate, of distinct P0 stromal subpopulations. Such a culture-induced intermediate has previously been reported for joint fibroblasts (Wei et al., 2020). These stark changes in subset identity are consistent with phenotypic changes across cell culture passage reported by bulk sequencing (Machaliński et al., 2020) and flow cytometry (Philippeos et al., 2018). However, our work describing fibrosis-related cell behavior of cell and ECM alignment, which is remarkably persistent through passages in culture (Kenny et al, 2023), indicates aspects of disease-relevant cell biology can be modelled in standard culture. This highlights the importance of understanding the strengths/weaknesses of models being used and value in developing strategies to recapitulate specific *in vivo* cell features.

The strength of scRNA-seq is to characterize the presence and proportions of cell subtypes. Here, we confirmed mesenchymal fibroblasts are abnormally enriched in keloid scars versus normal skin. This is consistent with the original individual reports (Deng et al., 2021, Liu et al., 2021), and supports a possible role for these cells in the pathology. Building on decades of studies showing that fibroblasts isolated from keloid and other fibrotic tissues are transcriptionally distinct from controls, we also aimed to explore disease-associated transcriptional differences within specific fibroblast subsets, which can be examined using pseudo-bulk approaches. Here we focused on the mesenchymal fibroblast population and detected hundreds of differentially expressed genes between keloid and normal skin mesenchymal fibroblasts. Yet, most of these differences did not persist in culture. This finding indicates that comparing cells using standard *in vitro* conditions may miss and/or underestimate biological differences within certain cell subsets. However, we still detected 295 differential genes in cultured keloid cells, indicating many disease-relevant characteristics persist or emerge. In particular, SOCS3, a known STAT-response gene and downstream target of IL-6 signaling, was upregulated in keloids at P8/P9, which further supports our previous data that autocrine IL-6 signaling is uniquely active in cultured keloid fibroblasts (Kenny et al., 2023). Additional studies powered to detect within-cell-type, disease-associated transcriptional differences are needed. Future work could address what features of the culture system (e.g. serum (Iyer et al., 1999), substrate stiffness (Ross et al., 2024, Son et al., 2024), absence/presence of specific supplements) are particularly influential to inducing differential expression at later passages, and indeed, whether they equally affect healthy and keloid fibroblasts.

With an aim to develop an *in vitro* model to investigate the mesenchymal cellular phenotype in keloids, strategies to maintain this fibroblast subset were explored. Others have reported that positional signals can retain fibroblast subtype identity; for example, Wei et al (2020) showed that synovial sublining versus lining fibroblasts are maintained by NOTCH signaling from nearby endothelial cells. We used cell-cell communication analysis to identify heavily utilized incoming signaling pathways. Periostin was identified as an autocrine signal for mesenchymal fibroblasts, consistent with previous reports (Supplementary Figure 3) (Deng et al., 2021). We particularly noted periostin as an important incoming signal for P4 CEMIP^+^ fibroblasts, yet the main source is the SRGN^+^ population, which at this passage is much-reduced (Supplementary Figure 4). Along with other ECM interaction pathways (collagen, fibronectin), this indicated that keloid-relevant mesenchymal fibroblasts might be retained by the presence of ECM in the culture conditions. We tested this using ascorbic acid, an important cofactor needed for collagen secretion and the stabilization of other ECM components (Kao et al., 1990, Phillips et al., 1994). As hypothesized, adding ascorbic acid into standard culture conditions for just 5 days resulted in a transcriptional shift towards a mesenchymal phenotype. Similarly, transitions towards more *in vivo*-like phenotypes with cultures inclusive of skin ECM derivatives have been described previously (Zhang et al., 2009), but this has not included in-depth transcriptional analysis of individual subset markers. Furthermore, the powerful transcriptional effects of the ECM have been demonstrated previously with mesenchymal stem cells, with differing composition leading to distinct cell lineages (Cai et al., 2015, Hoshiba et al., 2009, 2010, Ozguldez et al., 2018). Though the cell-derived ECM in this study was not characterized, future studies could evaluate whether keloid-specific ECM components (Barallobre-Barreiro et al., 2019, Zhang et al., 2021) are stably expressed past P4 and if they may have specific effects on sustaining keloid cell phenotypes.

There are now many commercially available media formulations specific for primary fibroblasts (e.g. FGM-2, FibroGro; some supplemented with FGF, insulin) that may alter these findings again; our data may serve as a resource, highlighting marker genes for characterizing cells resulting from different expansion conditions. Here we use the ECM as an example of a wide-ranging incoming signal that could potentially enrich or maintain mesenchymal fibroblasts, but it is hoped that the findings can also be exploited for other fibroblast subsets, e.g., NEGR (neuronal growth regulator) manipulation may uniquely promote a proinflammatory phenotype (Figure 6b).

In conclusion, this study describes the specific fibroblast subset changes that occur after multiple passages, and the impact this has for investigating pathological mechanisms in the context of keloid fibroblasts. Furthermore, we describe a potential method for adjusting culture conditions in an attempt to reverse loss of specific phenotypes. As the field works towards increasingly sophisticated *in vitro* models of disease (e.g. 3D, complex organoids), knowledge of how culture components may affect cell identity is important. Future work dedicated to establishing protocols to isolate and preserve distinct subsets will be valuable, as accurate models are required for scientific understanding and the development of much-needed novel therapeutics.

## MATERIALS AND METHODS

### Ethics and patient samples

All samples were collected under research ethics committee (REC) reference: 14/NS/1073. See Supplementary Table 1 for donor demographics.

### Tissue dissociation

Tissue samples were washed successively in 10% w/w iodinated povidone (0.025g/10mL, Ecolab, #3030440), 70% ethanol (Sigma Aldrich, #32221) and twice in 0.05mg/mL gentamycin solution (Sigma Aldrich, #G1397), before being soaked in 10mg/mL Dispase II solution (Gibco, #17105041) overnight at 4°C. Following epidermis removal, the remaining dermis was dissociated using a Human Whole Skin Dissociation kit (Miltenyi Biotec, #130-101-540) as per manufacturer’s instructions. The resulting cell suspension was frozen at -80°C in complete media with 40% fetal bovine serum (FBS; Cytiva, #SV30160.03) and 10% DMSO (Sigma Aldrich, #D2650) before being moved to vapour-phase nitrogen until culture.

### Cell culture

#### General fibroblast culture

Normal and keloid dermal cells/fibroblasts were defrosted and grown until 80% confluency in 2D in ‘complete medium’: high-glucose Dulbecco’s Modified Eagle Medium (DMEM with GlutaMAX and pyruvate, Gibco, #31966021) + 10% FBS (Cytiva, #SV30160.03) and 1% penicillin/streptomycin (Sigma-Aldrich, #P0781). Cells were passaged according to standard protocols, detaching them by incubation with trypsin-EDTA solution (Sigma-Aldrich, #T4174) for 5 mins at 37°C. For P0/P4 scRNA-seq, P0 vials of normal and keloid dermal cells were thawed and then cultured in complete medium over four passages. These cells (P4) were then frozen at - 80°C in complete medium with 40% FBS and 10% DMSO before being moved to vapour-phase nitrogen. Immediately prior to sequencing, both P0 and P4 vials were thawed and enriched for live cells using the MACS Dead Cell Removal Kit (Miltenyi Biotec, #130-090-101).

For qPCR experiments, cultures were lysed in 350μL RLT buffer (Qiagen, #74104), supplemented with 1% beta-mercaptoethanol (PanReac AppliChem, #A1108.0100) before being frozen at -80°C until downstream analysis.

#### Fibroblast culture with cell-derived matrices

To study the effects of matrix deposition on fibroblast phenotype, a protocol based on cell-derived matrices was adapted and utilised from (Kaukonen et al., 2017). First, sterile 13 mm glass coverslips (VWR, #631-1578) were added to a 24 well plate (VWR, #734-1605) and rinsed in 70% ethanol for at least 15 minutes. After PBS washing, sterile 0.2% (wt/vol) gelatin (from bovine skin, Sigma, #G9391) was added for 60 minutes at 37°C. Coverslips were then rinsed and the gelatin crosslinked with 1% glutaraldehyde (Alfa Aesar, #A10500.22) for 30 minutes at room temperature. Coverslips were rinsed with sterile PBS and incubated with 1M Glycine (Sigma, #G8790) for 20 minutes at room temperature to quench crosslinking. Coverslips were rinsed with PBS and incubated with complete medium for 30 minutes at 37°C and either used immediately, or were filled with at least 1mL of PBS and stored for up to 1 month at 4°C.

For bulk RNA sequencing, normal and keloid dermal fibroblasts (P8-P9) were thawed and seeded in triplicate at a density of 26,300 cells/cm² (24-well plate, near confluency), and left overnight to adhere and spread. Cells were cultured in complete medium with or without supplementation with 50μg/mL ascorbic acid (Sigma, #A4403) for 5 days, with media changes every other day. After 5 days, triplicate wells were combined into 350μL RLT buffer (Qiagen, #74104), supplemented with 1% beta-mercaptoethanol (PanReac AppliChem, #A1108.0100) before being frozen at -80°C until downstream analysis. This experiment was repeated on serial passages, P8 – P9, which were used as technical replicates (N=4).

### Single cell RNA sequencing

#### Library preparation and sequencing

RNA from P0/P4 matched samples was given to the Waterloo Genomics Centre, King’s College London for generation of single-cell gel beads, barcoding, clean up, cDNA amplification and library construction with a Chromium Single Cell 3’ Reagent Kit v3.1 (10X Genomics). The resulting libraries were sequenced on an Illumina HiSeq 2000 platform (Illumina) by Novogene (Novogene Co. Ltd, Cambridge, UK).

#### Data analysis

Raw sequencing fastq files were processed using the kb-python package (Melsted et al., 2021). Briefly, a pseudoalignment index was created using the *Homo sapiens* Ensembl reference transcriptome (GRCh38.p14). Read matrices were converted to Seurat objects using the Seurat package (v4.3.0) (Butler et al., 2018, Satija et al., 2015, Stuart et al., 2019), allowing for quality control (QC) and analysis (authors’ standard workflow); QC filters applied: nFeature_RNA <5000 & >500, percent.mt <15. All subsequent analyses were performed using Seurat R package version 4.3.0, with detailed scripts provided in ’Supplementary_Script’. Processed and raw data are available under GEO accession GSE293834, with the following Seurat objects: all P0/P4 data before (GSE293834_processed.Robj.gz) and after SCTransform (GSE293834_sct_v4.Robj.gz); the stromal cell subsets only (GSE293834_sct_v4_stromal.Robj.gz) as well as the P4 only subset (GSE293834_P4_only.Robj.gz). See also Supplementary Table 3 for normalization parameters. Clusters emerging from the unsupervised analysis were assigned cell types by cross-referencing the top 15 DEGs (by proportion) with the Panglao (Franzén et al., 2019) and Human Protein Atlas (Regev et al., 2017) online databases. The P0/P4 dataset did not require integration, as samples were run on the same 10X chip in batch-controlled fashion. Previously published keloid scRNA-seq datasets (Deng et al., 2021, Direder et al., 2022a, Liu et al., 2021) were computationally integrated in Seurat by identifying variable features across individual biopsies (Supplementary Figure 5). Finally, the CellChat package (Jin et al., 2021) (v 1.5.0) was used to analyse the integrated dataset, following the authors’ vignette.

To identify a keloid mesenchymal fibroblast transcriptional phenotype, pseudobulk counts were generated from the integrated scRNA-seq stromal-only Seurat object, which was further subsetted to include only mesenchymal fibroblasts from keloid and normal skin samples (excluding normal scar). Pseudobulk counts were generated by first normalising counts using the ‘RC’ method (Seurat V4) and then summing across all cells per sample. Differential gene expression analysis was performed in R using DESeq2 (Love et al., 2014), and visualised with volcano plots generated using the EnhancedVolcano package (Blighe et al., 2019). The resulting gene list was then filtered for expressed genes (basemean>50) and for significant (padj<0.05) upregulated (Log2FC>0.5) and downregulated (Log2FC<-0.5) genes.

### RNA extraction and quantification

RNA was purified using either the RNeasy Mini kit (for bulk RNA sequencing) (Qiagen, #74104) or the RNeasy Micro Plus kit (for RT-qPCR) (Qiagen, #74034) following manufacturer’s instructions, and quantified using a Qubit high sensitivity RNA assay (Invitrogen, #Q32852). RNA quality was ascertained by either Bioanalyzer 2100 (Agilent) (for bulk RNA sequencing) or Tapestation 4150 (Agilent) (for RT-qPCR). In both cases, RNA Integrity Number (RIN) values above 9 were used.

### Bulk RNA sequencing

#### Sample preparation

RNA from P8-9 normal and keloid fibroblasts cultured with and without ascorbic acid were shipped to BGI Genomics Co. Ltd., Hong Kong for library preparation and sequencing using the BGISEQ platform. The SOAPnuke software was used to filter out low-quality reads, with remaining clean reads aligned to the reference human genome (Homo Sapiens GRCh37) and using HISAT2 and Bowtie2 software. All fastq and processed files are available on the GEO repository under accession number GSE303591. Transcripts per million (TPM) values were used for analyses in Figures 4a-b and 6d-h. For comparison to pseudobulk differential expression in Figure 5c, raw read counts from our bulk-RNA seq were normalised using the same method as DESeq2 (estimateSizeFactors) and then filtered for truly expressed genes (TPM>1 in all samples in normal skin or keloid group).

#### Analysis

##### Comparison to ex vivo keloid mesenchymal fibroblast differential expression

To evaluate whether P8-9 keloid fibroblasts maintained *ex vivo* keloid mesenchymal differential gene expression, Log_2_FC values from the bulk sequenced keloid control (i.e. absence of ascorbic acid) samples were compared to the pseudobulk-DESeq2 outputs from keloid versus normal skin mesenchymal fibroblasts. Log_2_FC values for each bulk RNA sequenced keloid control sample was calculated based on comparison to the mean average normalised read count value of the normal control group.

##### Gene set variation analysis (GSVA)

To evaluate the change in cell phenotype with and without ascorbic acid treatment, gene set variation analysis (GSVA) was performed using gene signatures of *ex vivo* fibroblast subsets derived from the integrated scRNA sequencing data (Deng et al., 2021, Direder et al., 2022a, Liu et al., 2021). Signatures for each cluster were compiled based on genes that were expressed at Log_2_FC>1.5 or ratio>1.5 (padj<0.05) compared to all other stromal subsets (Supplementary Data 1). Log TPM values were used for analysis to avoid skewing by extreme values. Analysis was carried out using the GSVA R package (Hänzelmann et al., 2013).

### RT-qPCR

RNA was reverse transcribed using SuperScript III reverse transcriptase (200U/μL) with dithiothreitol (0.1M) and first-strand buffer (5X) (Invitrogen, #18080093) and random primers (Invitrogen, #48190011) plus dNTPs (Invitrogen, #18427013) according to manufacturer’s instructions. Due to low RNA yield (<4ng/μL) from some samples, RNA from all samples for the P0-P4 sequencing validation experiment were amplified and converted to cDNA using the Smart-seq2 protocol (steps 1 to 27) (Picelli et al., 2014). cDNA was diluted to 2ng/μL, and qPCR was performed using a Roche LightCycler 480II with SYBR Green I (2x; Roche, #04887352001) according to manufacturer’s instructions. Primer sequences (tested for efficiency (80-120 accepted) and specificity (gel electrophoresis) are listed in Supplementary Table 2.

## Supporting information

CODE AVAILABILITY

CODE AVAILABILITY

## ABBREVIATIONS

scRNA-seq: Single-cell RNA sequencing
P0: non-passaged (not cultured)
P4: 4th passage
SMC: Smooth muscle cell

## DATA AVAILABILITY

Generated scRNA-seq and bulk RNA-seq data are available in NCBI’s Gene Expression Omnibus (GEO) and accessible through accession numbers GSE303591(bulk RNA-seq) and GSE293834 (scRNA-seq).

Acquired scRNA-seq data are accessible by the following:

Liu et al. (2021): <u>HRA000425</u>, hosted on the Genome Sequence Archive (GSA).

Deng et al. (2021): <u>GSE163973</u>, hosted in the GEO repository.

Direder et al. (2022a): <u>GSE181316</u>, hosted in the GEO repository.

## CODE AVAILABILITY

Code to reproduce the analyses described in this manuscript can be found in the Supplementary Script Files (*.html and *.rmd provided).

## ACKNOWLEDGEMENTS

AL was supported by the UK Medical Research Council (MR/N013700/1) and King’s College London as a member of the MRC Doctoral Training Partnership in Biomedical Sciences. This work was further supported by the MRC IAA 2021 King’s College London (MR/X502923/1; FD, TS, AL) and the Wellcome Trust (107859/Z/15/Z; BS).

ED was supported by King’s College London as part of the Cell Therapies & Regenerative Medicine Wellcome Trust PhD Programme (108874/Z/15/Z).

For the purpose of Open Access, the author has applied a CC BY public copyright licence to any Author Accepted Manuscript (AAM) version arising from this submission.

## AUTHOR CONTRIBUTIONS

Conceptualization: AL, BS, TS, FD; Data Curation: AL; Formal Analysis: AL; Funding Acquisition: TS, FD; Investigation: AL; Methodology: AL, DF, TS, FD; Project Administration: TS, FD; Resources: AL, TS, FD, DF; Software: AL; Supervision: TS, FD; Validation: AL, TS, FD; Visualization: AL; Writing - Original Draft Preparation: AL; Writing - Review and Editing: AL, TS, FD, DF, BS.

## REFERENCES

Barallobre-Barreiro J, Woods E, Bell RE, Easton JA, Hobbs C, Eager M, et al. Cartilage-like composition of keloid scar extracellular matrix suggests fibroblast mis-differentiation in disease. Matrix Biology Plus 2019;4:100016.

Blighe K, Rana S, Lewis M. EnhancedVolcano: Publication-ready volcano plots with enhanced colouring and labeling. R package version 2019;1(0):10.18129.

Buechler MB, Pradhan RN, Krishnamurty AT, Cox C, Calviello AK, Wang AW, et al. Cross-tissue organization of the fibroblast lineage. Nature 2021;593(7860):575–9.

Butler A, Hoffman P, Smibert P, Papalexi E, Satija R. Integrating single-cell transcriptomic data across different conditions, technologies, and species. Nat Biotechnol 2018;36(5):411–20.

Cai R, Nakamoto T, Kawazoe N, Chen G. Influence of stepwise chondrogenesis-mimicking 3D extracellular matrix on chondrogenic differentiation of mesenchymal stem cells. Biomaterials 2015;52:199–207.

Chike-Obi CJ, Cole PD, Brissett AE. Keloids: pathogenesis, clinical features, and management. Semin Plast Surg 2009;23(3):178–84.

Deng C-C, Hu Y-F, Zhu D-H, Cheng Q, Gu J-J, Feng Q-L, et al. Single-cell RNA-seq reveals fibroblast heterogeneity and increased mesenchymal fibroblasts in human fibrotic skin diseases. Nature Communications 2021;12(1):3709.

Dirand Z, Tissot M, Chatelain B, Viennet C, Rolin G. Is Spheroid a Relevant Model to Address Fibrogenesis in Keloid Research? Biomedicines 2023;11(9):2350.

Direder M, Weiss T, Copic D, Vorstandlechner V, Laggner M, Pfisterer K, et al. Schwann cells contribute to keloid formation. Matrix Biol 2022a;108:55–76.

Direder M, Wielscher M, Weiss T, Laggner M, Copic D, Klas K, et al. The transcriptional profile of keloidal Schwann cells. Exp Mol Med 2022b;54(11):1886–900.

Franzén O, Gan L-M, Björkegren JLM. PanglaoDB: a web server for exploration of mouse and human single-cell RNA sequencing data. Database 2019;2019.

Friedman DW, Boyd CD, Mackenzie JW, Norton P, Olson RM, Deak SB. Regulation of collagen gene expression in keloids and hypertrophic scars. J Surg Res 1993;55(2):214–22.

Hänzelmann S, Castelo R, Guinney J. GSVA: gene set variation analysis for microarray and RNA-Seq data. BMC Bioinformatics 2013;14(1):7.

Harper RA. Keloid fibroblasts in culture: Abnormal growth behaviour and altered response to the epidermal growth factor. Cell Biology International Reports 1989;13(4):325–35.

Hoshiba T, Kawazoe N, Tateishi T, Chen G. Development of Stepwise Osteogenesis-mimicking Matrices for the Regulation of Mesenchymal Stem Cell Functions*. Journal of Biological Chemistry 2009;284(45):31164–73.

Hoshiba T, Kawazoe N, Tateishi T, Chen G. Development of Extracellular Matrices Mimicking Stepwise Adipogenesis of Mesenchymal Stem Cells. Advanced Materials 2010;22(28):3042–7.

Iyer VR, Eisen MB, Ross DT, Schuler G, Moore T, Lee JCF, et al. The Transcriptional Program in the Response of Human Fibroblasts to Serum. Science 1999;283(5398):83–7.

Jin S, Guerrero-Juarez CF, Zhang L, Chang I, Ramos R, Kuan C-H, et al. Inference and analysis of cell-cell communication using CellChat. Nature Communications 2021;12(1):1088.

Kao J, Huey G, Kao R, Stern R. Ascorbic acid stimulates production of glycosaminoglycans in cultured fibroblasts. Experimental and Molecular Pathology 1990;53(1):1–10.

Kaukonen R, Jacquemet G, Hamidi H, Ivaska J. Cell-derived matrices for studying cell proliferation and directional migration in a complex 3D microenvironment. Nat Protoc 2017;12(11):2376–90.

Kenny FN, Marcotti S, De Freitas DB, Drudi EM, Leech V, Bell RE, et al. Autocrine IL-6 drives cell and extracellular matrix anisotropy in scar fibroblasts. Matrix Biology 2023;123:1–16.

Kim M, Mirsky N, Spielman A, Mathew P, Yechieli R, Tang JC, et al. Evaluating Symptomatic and Psychosocial Well-being After Keloid Treatment With SCAR-Q. Aesthet Surg J 2022;42(6):Np416-np22.

Korsunsky I, Wei K, Pohin M, Kim EY, Barone F, Major T, et al. Cross-tissue, single-cell stromal atlas identifies shared pathological fibroblast phenotypes in four chronic inflammatory diseases. Med 2022;3(7):481–518.e14.

Ladin DA, Hou Z, Patel D, McPhail M, Olson JC, Saed GM, et al. p53 and apoptosis alterations in keloids and keloid fibroblasts. Wound Repair Regen 1998;6(1):28–37.

Liu X, Chen W, Zeng Q, Ma B, Li Z, Meng T, et al. Single-cell RNA-seq reveals lineage-specific regulatory changes of fibroblasts and vascular endothelial cells in keloids. J Invest Dermatol 2021.

Love MI, Huber W, Anders S. Moderated estimation of fold change and dispersion for RNA-seq data with DESeq2. Genome Biol 2014;15(12):550.

Machaliński B, Rogińska D, Szumilas K, Zawiślak A, Wilk A, Stecewicz I, et al. Transcriptome Profile of Human Fibroblasts in an Ex Vivo Culture. Int J Med Sci 2020;17(1):125–36.

Melsted P, Booeshaghi AS, Liu L, Gao F, Lu L, Min KH, et al. Modular, efficient and constant-memory single-cell RNA-seq preprocessing. Nature Biotechnology 2021;39(7):813–8.

Neumann E, Riepl B, Knedla A, Lefèvre S, Tarner IH, Grifka J, et al. Cell culture and passaging alters gene expression pattern and proliferation rate in rheumatoid arthritis synovial fibroblasts. Arthritis Research & Therapy 2010;12(3):R83.

Ozguldez HO, Cha J, Hong Y, Koh I, Kim P. Nanoengineered, cell-derived extracellular matrix influences ECM-related gene expression of mesenchymal stem cells. Biomaterials Research 2018;22(1):32.

Philippeos C, Telerman SB, Oulès B, Pisco AO, Shaw TJ, Elgueta R, et al. Spatial and Single-Cell Transcriptional Profiling Identifies Functionally Distinct Human Dermal Fibroblast Subpopulations. J Invest Dermatol 2018;138(4):811–25.

Phillips CL, Combs SB, Pinnell SR. Effects of ascorbic acid on proliferation and collagen synthesis in relation to the donor age of human dermal fibroblasts. J Invest Dermatol 1994;103(2):228–32.

Picelli S, Faridani OR, Björklund ÅK, Winberg G, Sagasser S, Sandberg R. Full-length RNA-seq from single cells using Smart-seq2. Nature Protocols 2014;9(1):171–81.

Regev A, Teichmann SA, Lander ES, Amit I, Benoist C, Birney E, et al. The Human Cell Atlas. Elife 2017;6.

Ross R, Guo Y, Walker RN, Bergamaschi D, Shaw TJ, Connelly JT. Biomechanical Activation of Keloid Fibroblasts Promotes Lysosomal Remodeling and Exocytosis. Journal of Investigative Dermatology 2024;144(12):2730–41.

Satija R, Farrell JA, Gennert D, Schier AF, Regev A. Spatial reconstruction of single-cell gene expression data. Nature Biotechnology 2015;33(5):495–502.

Solé-Boldo L, Raddatz G, Schütz S, Mallm JP, Rippe K, Lonsdorf AS, et al. Single-cell transcriptomes of the human skin reveal age-related loss of fibroblast priming. Commun Biol 2020;3(1):188.

Son DO, Benitez R, Diao L, Hinz B. How to Keep Myofibroblasts under Control: Culture of Mouse Skin Fibroblasts on Soft Substrates. Journal of Investigative Dermatology 2024;144(9):1923–34.

Stuart T, Butler A, Hoffman P, Hafemeister C, Papalexi E, Mauck WM, et al. Comprehensive Integration of Single-Cell Data. Cell 2019;177(7):1888–902.e21.

van Vijven M, Wunderli SL, Ito K, Snedeker JG, Foolen J. Serum deprivation limits loss and promotes recovery of tenogenic phenotype in tendon cell culture systems. Journal of Orthopaedic Research 2021;39(7):1561–71.

Wei K, Korsunsky I, Marshall JL, Gao A, Watts GFM, Major T, et al. Notch signalling drives synovial fibroblast identity and arthritis pathology. Nature 2020;582(7811):259–64.

Yang Y-HK, Ogando CR, Wang See C, Chang T-Y, Barabino GA. Changes in phenotype and differentiation potential of human mesenchymal stem cells aging in vitro. Stem Cell Research & Therapy 2018;9(1):131.

Zhang S, Liu B, Wang W, Lv L, Gao D, Chai M, et al. The “Matrisome” reveals the characterization of skin keloid microenvironment. The FASEB Journal 2021;35(4):e21237.

Zhang Y, He Y, Bharadwaj S, Hammam N, Carnagey K, Myers R, et al. Tissue-specific extracellular matrix coatings for the promotion of cell proliferation and maintenance of cell phenotype. Biomaterials 2009;30(23-24):4021–8.

