## Supplementary Figure for "Single-cell RNA sequencing clarifies dermal fibroblast subset representation *in vitro* and reveals variable persistence of keloid disease-associated features"

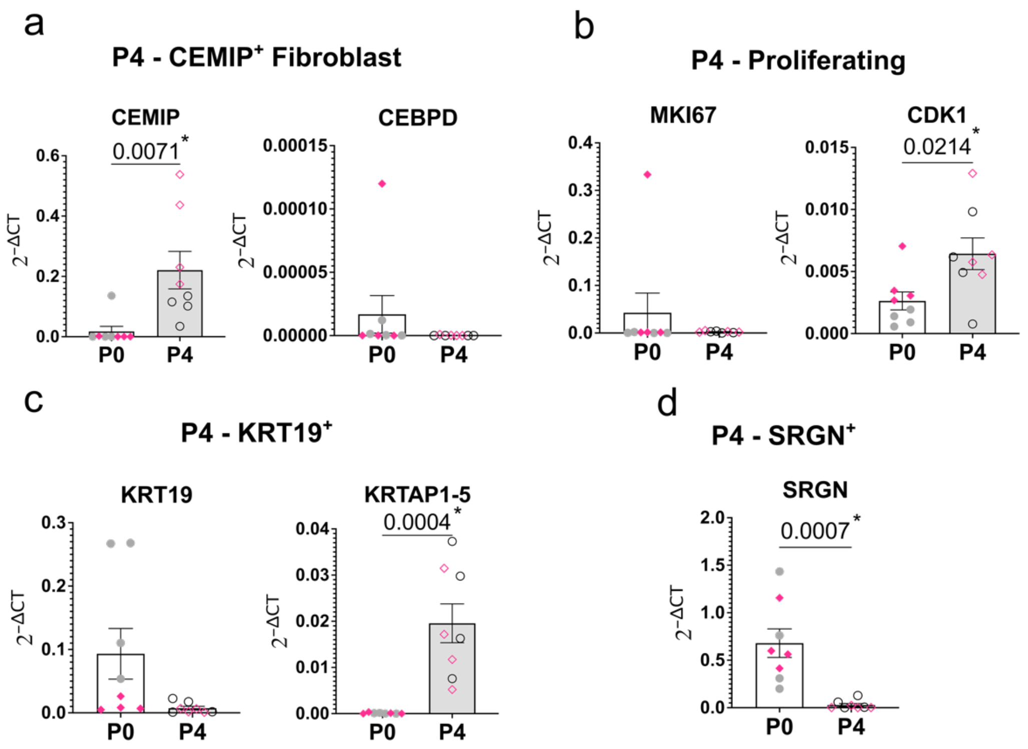


**Supplementary Figure 1. Confirmation of P4 cell subsets in further expanded dermal cell samples.**

(a-d) Expression of passage (P) 4 subset markers in P0 (white bar) and P4 (grey bar) normal (grey circles) and keloid (pink diamonds) dermal cells. N=8/group, with each dot derived from a different donor. qPCR expression normalized to reference gene, GAPDH. Bars represent mean +/- SEM. Statistical test: Unpaired T-test; p-values are displayed on the graph.

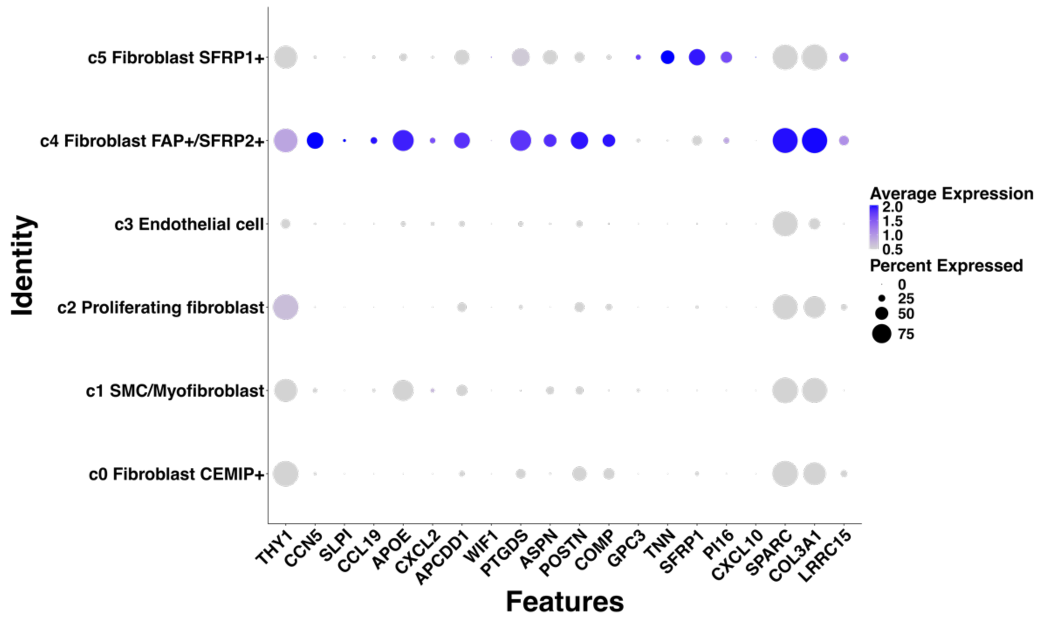


**Supplementary Figure 2. Expression of subset specific and universal fibroblast markers in P0/P4 stromal cells.**

Dot plot showing the expression of marker genes across each stromal cluster in the P0/P4 dataset. Blue color gradient represents average expression of given gene. Circle size represents the percentage of cells within a cluster expressing a given gene.

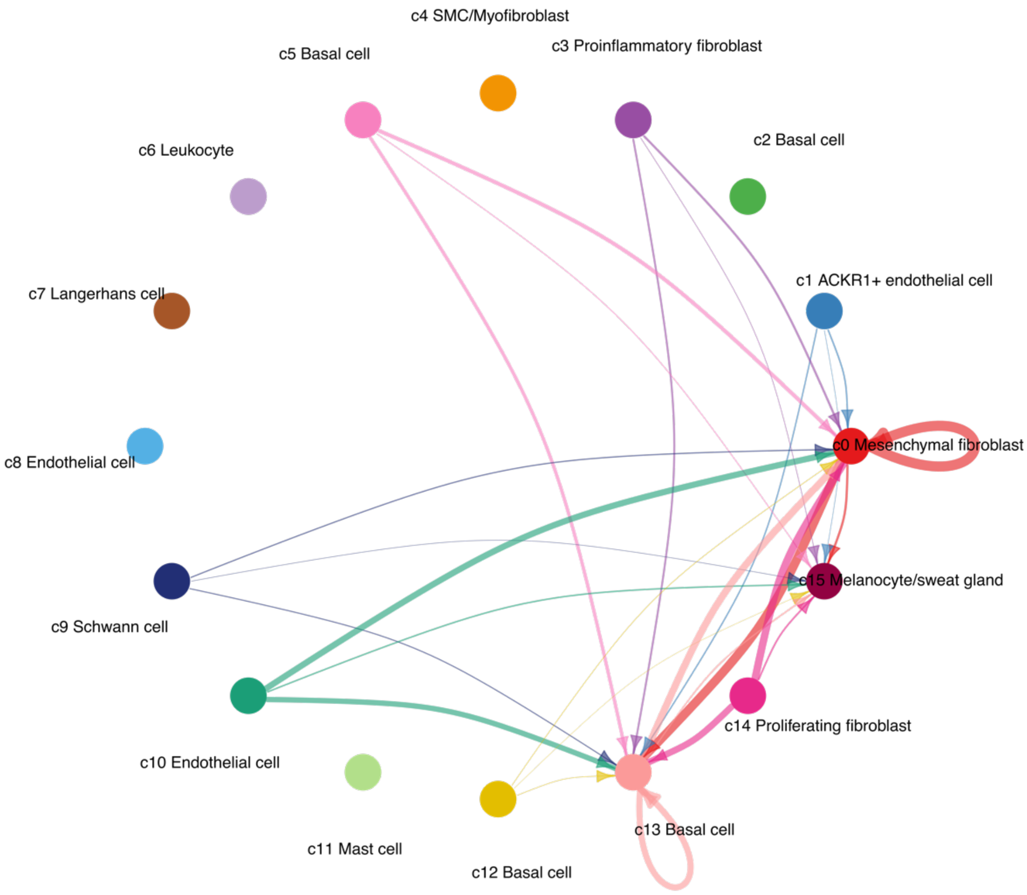


**Supplementary Figure 3. Mesenchymal fibroblasts may receive periostin signaling from several cell types and may act in an autocrine signaling loop.**

Analysis using data derived from integrating previously published scRNA-seq studies of keloid and normal skin/scar (see Methods). Circle plot illustrating the periostin signaling network (POSTN-ITGAV:ITGB3). Arrowheads point toward receivers (direction of signaling input). The thickness of lines represents the strength of signal. Plot created using the package CellChat.

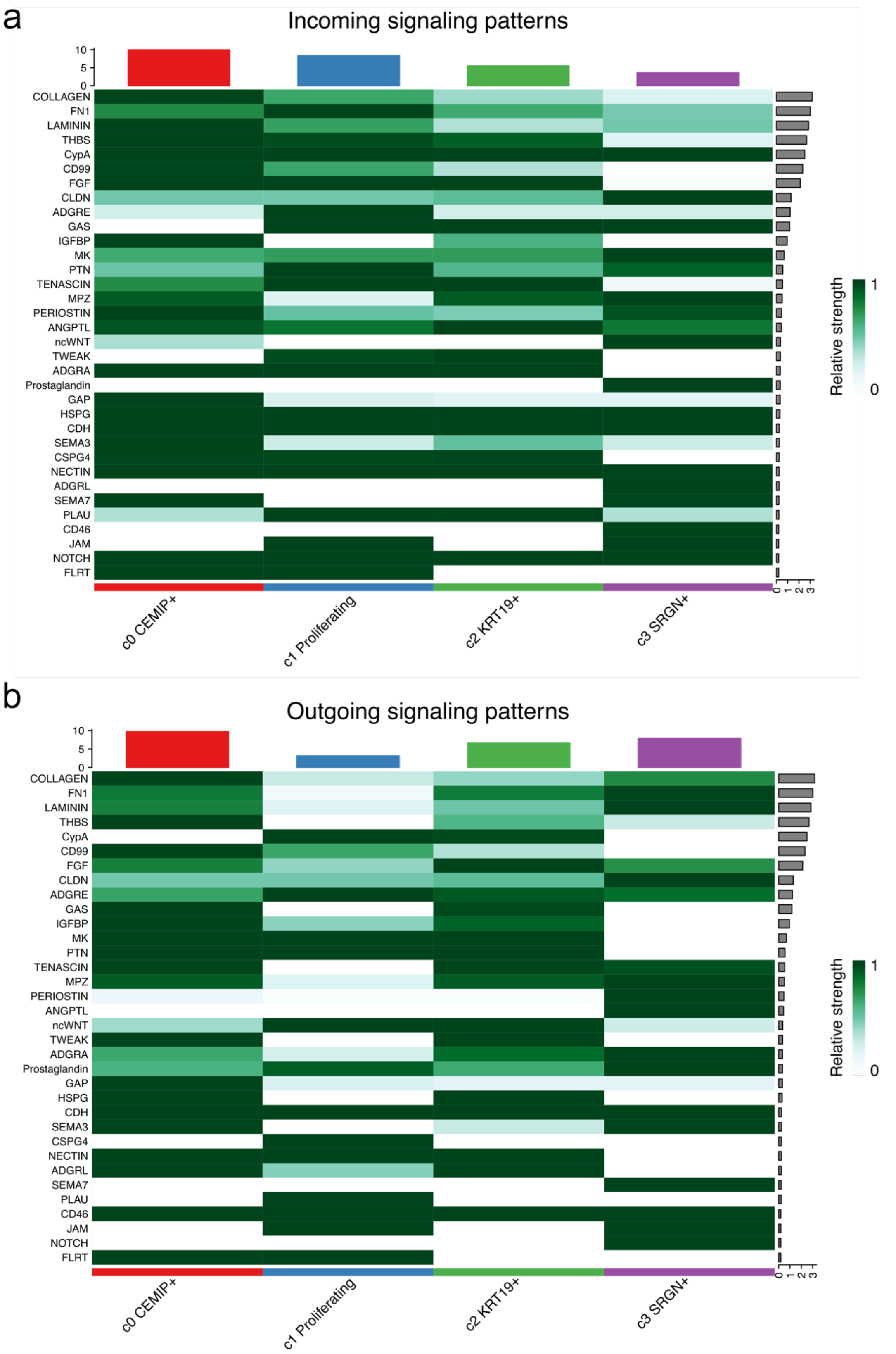


**Supplementary Figure 4. Cell-cell communication analysis highlights potential pathways preserved at later passage culture.**

(a-b) Cell-cell communication analysis of (a) incoming and (b) outgoing signals within P4-only cell clusters. The green color bar represents the relative signaling strength of pathways across cell subsets, with darker green indicating higher strength (row-scaled values). The colored bar plot at the top shows the total signaling strength of each cell subset, summarizing all pathways. The right grey bar plot indicates the total signaling strength of each pathway, summarizing all cell subsets. Generated using the CellChat package.

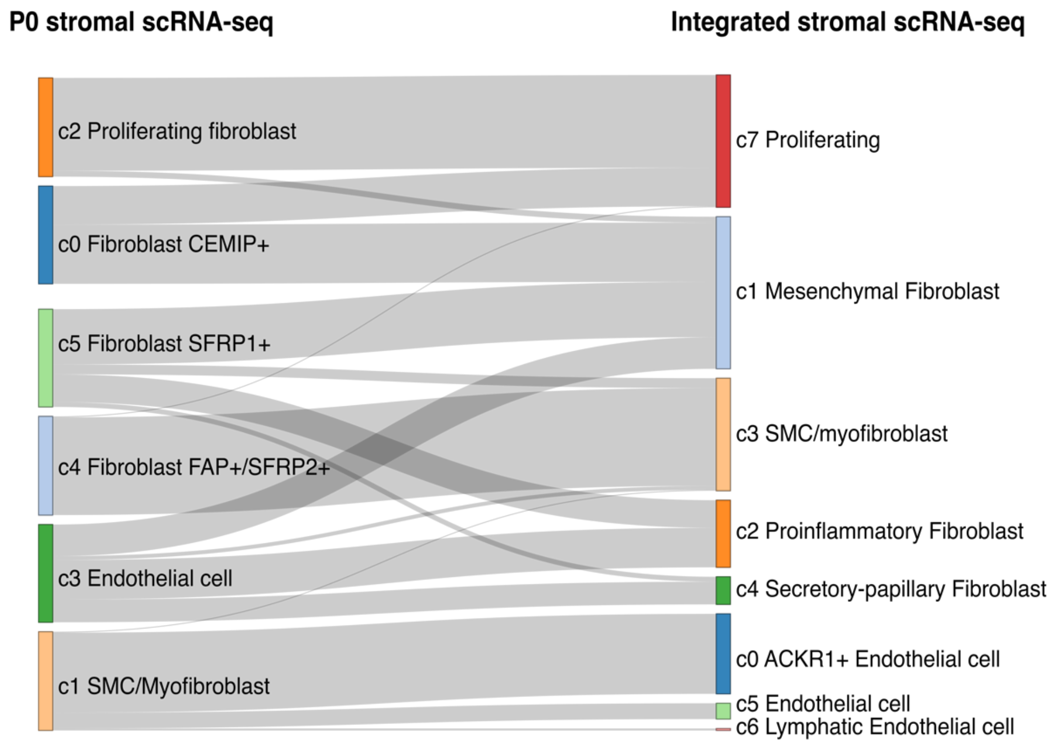


**Supplementary Figure 5. Mapping of P0 annotated stromal subsets to integrated scRNA-seq dataset.**

Sankey plot illustrates how P0/P4 stromal subsets identified in this study map to the integrated stromal dataset clusters. The thickness of lines is proportional to the number of cells.
