## Supplementary Table for "Single-cell RNA sequencing clarifies dermal fibroblast subset representation *in vitro* and reveals variable persistence of keloid disease-associated features"

**Supplementary Table 1. Donor demographics**

| Sample ID | Normal/Keloid | Gender | Age at excision | Scar location | Passage |
|---|---|---|---|---|---|
| P0/P4 scRNA-seq and P8/P9 bulk RNAseq (Figure 1-3, Figure 4a-b, Figure 5c, Figure 6d-h) | | | | | |
| 240216 | Normal & Keloid (paired) | M | 53 | Pelvis | 0 & 4<br>8 & 9 |
| P0/P4 Normal skin/Keloid scar samples (Figure 4c-f - qPCR validation) | | | | | |
| 280616B | Keloid | F | UNKNOWN | Ear | 0 |
| 170215 | Keloid | F | 24 | Ear | 0 & 4 |
| 180216 | Keloid | F | 50 | Abdomen | 0 |
| 090316 | Keloid | F | 50 | Umbilicus | 0 |
| 011015A | Keloid | M | 38 | Back | 4 |
| 240915 | Keloid | UNKNOWN | UNKNOWN | UNKNOWN | 4 |
| 181111 | Keloid | UNKNOWN | UNKNOWN | UNKNOWN | 4 |
| 090915A | Normal | F | 19 | Breast | 0 |
| 251115 | Normal | UNKNOWN | UNKNOWN | UNKNOWN | 0 & 4 |
| 160517 | Normal | UNKNOWN | UNKNOWN | Thigh | 0 & 4 |
| 300915 | Normal | F | 44 | Back | 0 & 4 |
| 020915 | Normal | F | 22 | Breast | 4 |

**Supplementary Table 2. Primers**

| Gene | Forward | Reverse |
|---|---|---|
| GAPDH | GGAAGGTGAAGGTCGGAGTCAAC | CAGAGTTAAAAGCAGCCCTGGT |
| ACKR1/DARC | AAGGATGGTCTTCTCATCTG | TTTTCACAAAGGCAGTGTAG |
| CDK1 | ATGAGGTAGTAACACTCTGG | CCTATACTCCAAATGTCAACTG |
| CEMIP | ACCGAGCACATTCCAACTACCG | GGCAGAGATGATTGAGAGGAACG |
| CEBPD | CAGACTTTTCAGACAAACCC | TTTCGATTTCAAATGCTGC |
| FAP | GAAGAGGAAATGCTTGCTAC | CTAGGATATTGTTCATCGCC |
| KRT19 | AACCATGAGGAGGAAATCAG | CATGACCTCATATTGGCTTC |
| KRTAP1-5 | AGTTCTCAGACTTTGCATTG | TTTGTAGCATTTCTGTGTCC |
| MKI67 | GACAGAGGTTCCTAAGAGAG | AACAATCAGATTTGCTTCCG |
| MYH11 | CTATCTGCTAGAAAAATCACGG | CACTTCTCATCTTCTCCTTG |
| SFRP1 | CTTAAGTGTGACAAGTTCCC | TTTTCATCCTCAGTGCAAAC |
| SFRP2 | GACCTAGACGAGACCATC | ATACCTTTGGAGCTTCCTC |
| SRGN | GAATCCTCAGTTCAAGGTTATC | GATCTTGTTGGATTCACCTG |
| STEAP4 | TGATTCATATGTGGCTTTGG | CAGTTTGGACTGGACAAATC |
| PECAM1 | TGGAAAGCAGATACTCTAGAACGG | GGGATGTGCATCTGGCCTT |

**Supplementary Table 3. scRNA-seq analysis parameters**

| Analysis | Prinicipal components used for RunUMAP | Resolution used for FindClusters | Normalisation |
|---|---|---|---|
| Liu et al dataset (Liu et al., 2021) – Overall clusters | 15 | 0.5 | Log |
| Deng et al (Deng et al., 2021) – Overall clusters | 15 | 0.4 | Log |
| Direder et al (Direder et al., 2022a) – Overall clusters | 20 | 0.3 | Log |
| Integrated dataset – Overall clusters | 20 | 0.2 | Log |
| Integrated dataset – Stromal clusters | 15 | 0.2 | Log |
| P0-P4 dataset - Overall clusters | 20 | 0.2 | SCTransform |
| P0-P4 dataset - Stromal clusters | 20 | 0.3 | SCTransform |
| P0-P4 dataset – P4 clusters | 20 | 0.15 | SCTransform |
