## Supplementary material for "Single-cell RNA sequencing clarifies dermal fibroblast subset representation *in vitro* and reveals variable persistence of keloid disease-associated features": CODE AVAILABILITY: SupplementaryScript_1.html

Supplementary\_R script


### Supplementary\_R script

###### Amy Lock

#### 2025-08-08

### Supplementary

#### Load libraries

```
#install.packages("devtools")  # if not already installed
#devtools::install_github("satijalab/seurat", ref = "v4.4.0")
library(Seurat)
```

```
## Attaching SeuratObject
```

```
## 'SeuratObject' was built under R 4.4.1 but the current version is
## 4.4.2; it is recomended that you reinstall 'SeuratObject' as the ABI
## for R may have changed
```

```
## Seurat v4 was just loaded with SeuratObject v5; disabling v5 assays and
## validation routines, and ensuring assays work in strict v3/v4
## compatibility mode
```

```
library(SeuratObject)
#remotes::install_github("samuel-marsh/scCustomize", ref = "v2.0.0")
library(scCustomize)
```

```
## scCustomize v2.0.0
## If you find the scCustomize useful please cite.
## See 'samuel-marsh.github.io/scCustomize/articles/FAQ.html' for citation info.
```

```
library(BUSpaRse)
library(ggplot2)
library(tidyverse)
```

```
## ── Attaching core tidyverse packages ──────────────────────── tidyverse 2.0.0 ──
## ✔ dplyr     1.1.4     ✔ readr     2.1.5
## ✔ forcats   1.0.0     ✔ stringr   1.5.1
## ✔ lubridate 1.9.4     ✔ tibble    3.3.0
## ✔ purrr     1.1.0     ✔ tidyr     1.3.1
```

```
## ── Conflicts ────────────────────────────────────────── tidyverse_conflicts() ──
## ✖ dplyr::filter() masks stats::filter()
## ✖ dplyr::lag()    masks stats::lag()
## ℹ Use the conflicted package (<http://conflicted.r-lib.org/>) to force all conflicts to become errors
```

```
library(DropletUtils)
```

```
## Loading required package: SingleCellExperiment
## Loading required package: SummarizedExperiment
## Loading required package: MatrixGenerics
## Loading required package: matrixStats
## 
## Attaching package: 'matrixStats'
## 
## The following object is masked from 'package:dplyr':
## 
##     count
## 
## 
## Attaching package: 'MatrixGenerics'
## 
## The following objects are masked from 'package:matrixStats':
## 
##     colAlls, colAnyNAs, colAnys, colAvgsPerRowSet, colCollapse,
##     colCounts, colCummaxs, colCummins, colCumprods, colCumsums,
##     colDiffs, colIQRDiffs, colIQRs, colLogSumExps, colMadDiffs,
##     colMads, colMaxs, colMeans2, colMedians, colMins, colOrderStats,
##     colProds, colQuantiles, colRanges, colRanks, colSdDiffs, colSds,
##     colSums2, colTabulates, colVarDiffs, colVars, colWeightedMads,
##     colWeightedMeans, colWeightedMedians, colWeightedSds,
##     colWeightedVars, rowAlls, rowAnyNAs, rowAnys, rowAvgsPerColSet,
##     rowCollapse, rowCounts, rowCummaxs, rowCummins, rowCumprods,
##     rowCumsums, rowDiffs, rowIQRDiffs, rowIQRs, rowLogSumExps,
##     rowMadDiffs, rowMads, rowMaxs, rowMeans2, rowMedians, rowMins,
##     rowOrderStats, rowProds, rowQuantiles, rowRanges, rowRanks,
##     rowSdDiffs, rowSds, rowSums2, rowTabulates, rowVarDiffs, rowVars,
##     rowWeightedMads, rowWeightedMeans, rowWeightedMedians,
##     rowWeightedSds, rowWeightedVars
## 
## Loading required package: GenomicRanges
## Loading required package: stats4
## Loading required package: BiocGenerics
## 
## Attaching package: 'BiocGenerics'
## 
## The following objects are masked from 'package:lubridate':
## 
##     intersect, setdiff, union
## 
## The following objects are masked from 'package:dplyr':
## 
##     combine, intersect, setdiff, union
## 
## The following object is masked from 'package:SeuratObject':
## 
##     intersect
## 
## The following objects are masked from 'package:stats':
## 
##     IQR, mad, sd, var, xtabs
## 
## The following objects are masked from 'package:base':
## 
##     anyDuplicated, aperm, append, as.data.frame, basename, cbind,
##     colnames, dirname, do.call, duplicated, eval, evalq, Filter, Find,
##     get, grep, grepl, intersect, is.unsorted, lapply, Map, mapply,
##     match, mget, order, paste, pmax, pmax.int, pmin, pmin.int,
##     Position, rank, rbind, Reduce, rownames, sapply, saveRDS, setdiff,
##     table, tapply, union, unique, unsplit, which.max, which.min
## 
## Loading required package: S4Vectors
## 
## Attaching package: 'S4Vectors'
## 
## The following objects are masked from 'package:lubridate':
## 
##     second, second<-
## 
## The following objects are masked from 'package:dplyr':
## 
##     first, rename
## 
## The following object is masked from 'package:tidyr':
## 
##     expand
## 
## The following object is masked from 'package:utils':
## 
##     findMatches
## 
## The following objects are masked from 'package:base':
## 
##     expand.grid, I, unname
## 
## Loading required package: IRanges
## 
## Attaching package: 'IRanges'
## 
## The following object is masked from 'package:lubridate':
## 
##     %within%
## 
## The following objects are masked from 'package:dplyr':
## 
##     collapse, desc, slice
## 
## The following object is masked from 'package:purrr':
## 
##     reduce
## 
## Loading required package: GenomeInfoDb
## Loading required package: Biobase
## Welcome to Bioconductor
## 
##     Vignettes contain introductory material; view with
##     'browseVignettes()'. To cite Bioconductor, see
##     'citation("Biobase")', and for packages 'citation("pkgname")'.
## 
## 
## Attaching package: 'Biobase'
## 
## The following object is masked from 'package:MatrixGenerics':
## 
##     rowMedians
## 
## The following objects are masked from 'package:matrixStats':
## 
##     anyMissing, rowMedians
## 
## 
## Attaching package: 'SummarizedExperiment'
## 
## The following object is masked from 'package:SeuratObject':
## 
##     Assays
## 
## The following object is masked from 'package:Seurat':
## 
##     Assays
```

```
library(devtools)
```

```
## Loading required package: usethis
```

```
library(glmGamPoi)
```

```
## 
## Attaching package: 'glmGamPoi'
## 
## The following object is masked from 'package:dplyr':
## 
##     vars
## 
## The following object is masked from 'package:ggplot2':
## 
##     vars
```

```
library(presto)
```

```
## Loading required package: Rcpp
## Loading required package: data.table
## 
## Attaching package: 'data.table'
## 
## The following object is masked from 'package:SummarizedExperiment':
## 
##     shift
## 
## The following object is masked from 'package:GenomicRanges':
## 
##     shift
## 
## The following object is masked from 'package:IRanges':
## 
##     shift
## 
## The following objects are masked from 'package:S4Vectors':
## 
##     first, second
## 
## The following objects are masked from 'package:lubridate':
## 
##     hour, isoweek, mday, minute, month, quarter, second, wday, week,
##     yday, year
## 
## The following objects are masked from 'package:dplyr':
## 
##     between, first, last
## 
## The following object is masked from 'package:purrr':
## 
##     transpose
```

```
library(UCell)
library(RColorBrewer)
library(cowplot)
```

```
## 
## Attaching package: 'cowplot'
## 
## The following object is masked from 'package:lubridate':
## 
##     stamp
```

```
library(patchwork)
```

```
## 
## Attaching package: 'patchwork'
## 
## The following object is masked from 'package:cowplot':
## 
##     align_plots
```

```
library(dplyr)
library(CellChat)
```

```
## Loading required package: igraph
## 
## Attaching package: 'igraph'
## 
## The following object is masked from 'package:GenomicRanges':
## 
##     union
## 
## The following object is masked from 'package:IRanges':
## 
##     union
## 
## The following object is masked from 'package:S4Vectors':
## 
##     union
## 
## The following objects are masked from 'package:BiocGenerics':
## 
##     normalize, path, union
## 
## The following objects are masked from 'package:lubridate':
## 
##     %--%, union
## 
## The following objects are masked from 'package:dplyr':
## 
##     as_data_frame, groups, union
## 
## The following objects are masked from 'package:purrr':
## 
##     compose, simplify
## 
## The following object is masked from 'package:tidyr':
## 
##     crossing
## 
## The following object is masked from 'package:tibble':
## 
##     as_data_frame
## 
## The following objects are masked from 'package:stats':
## 
##     decompose, spectrum
## 
## The following object is masked from 'package:base':
## 
##     union
```

```
options(future.globals.maxSize = 8 * 1024^3)
```

#### Create list of seurat objects from kb-Python outputs (one object per sample)

```
#list the output files

# Define parent directory where all sample folders are located
data_dir <- "/Users/amylock_soper/Desktop/Paper/RAW"

# List all sample folders
sample_folders <- list.dirs(data_dir, recursive = FALSE)
```

```
# Import the gene ID conversion table (once, since it's shared)
t2g <- read_tsv(file.path(data_dir, "t2g_human.txt"), col_names = FALSE, col_types = cols(.default = col_character()))
ens2sym <- t2g %>% dplyr::select(X2, X3) %>% distinct()
gene_map <- ens2sym %>% tibble::deframe()

# Create a list to store Seurat objects
seurat_list <- list()

# Loop over each sample
for (sample_path in sample_folders) {
  
  # Extract sample name from folder path
  sample_name <- basename(sample_path)
  
  # Define the path to the count data
  counts_path <- sample_path
  
  # Skip if the directory does not exist (avoids errors)
  if (!dir.exists(counts_path)) {
    print(paste("Skipping", sample_name, "- directory not found"))
    next
  }

  # Load count matrix
  res_mat <- read_count_output(counts_path, name = "cells_x_genes", tcc = FALSE)

  # Convert Ensembl gene IDs to gene symbols
  rownames(res_mat) <- gene_map[rownames(res_mat)] %>% unname()
  
  # Ensure unique rownames
  rownames(res_mat) <- make.unique(rownames(res_mat))

  # Create Seurat Object
  seu <- CreateSeuratObject(res_mat, project = sample_name, min.cells = 3)

  # Compute mitochondrial percentage
  seu[["percent.mt"]] <- PercentageFeatureSet(seu, pattern = "^MT-")

# Store processed Seurat object
  seurat_list[[sample_name]] <- seu

  # Print progress
  print(paste("Finished processing", sample_name, "| Cells remaining:", dim(seu)[2]))
}
```

```
## 'as(<dgTMatrix>, "dgCMatrix")' is deprecated.
## Use 'as(., "CsparseMatrix")' instead.
## See help("Deprecated") and help("Matrix-deprecated").
```

```
## Warning: Feature names cannot have underscores ('_'), replacing with dashes
## ('-')
## Warning: Feature names cannot have underscores ('_'), replacing with dashes
## ('-')
```

```
## Warning: The `slot` argument of `GetAssayData()` is deprecated as of SeuratObject 5.0.0.
## ℹ Please use the `layer` argument instead.
## ℹ The deprecated feature was likely used in the Seurat package.
##   Please report the issue at <https://github.com/satijalab/seurat/issues>.
## This warning is displayed once every 8 hours.
## Call `lifecycle::last_lifecycle_warnings()` to see where this warning was
## generated.
```

```
## [1] "Finished processing KDF0_counts_unfiltered | Cells remaining: 1419243"
```

```
## Warning: Feature names cannot have underscores ('_'), replacing with dashes
## ('-')
```

```
## Warning: Feature names cannot have underscores ('_'), replacing with dashes
## ('-')
```

```
## [1] "Finished processing KDF4_counts_unfiltered | Cells remaining: 1423066"
```

```
## Warning: Feature names cannot have underscores ('_'), replacing with dashes
## ('-')
## Warning: Feature names cannot have underscores ('_'), replacing with dashes
## ('-')
```

```
## [1] "Finished processing NDF0_counts_unfiltered | Cells remaining: 1341605"
```

```
## Warning: Feature names cannot have underscores ('_'), replacing with dashes
## ('-')
## Warning: Feature names cannot have underscores ('_'), replacing with dashes
## ('-')
```

```
## [1] "Finished processing NDF4_counts_unfiltered | Cells remaining: 1344420"
```

#### Unlist seurat objects and QC each individually

```
#renaming samples to shorter name
names(seurat_list) <- gsub("_counts_unfiltered$", "", names(seurat_list))  # Remove suffix
# Print new names to confirm
print(names(seurat_list))
```

```
## [1] "KDF0" "KDF4" "NDF0" "NDF4"
```

```
list2env(seurat_list, envir = .GlobalEnv)
```

```
## <environment: R_GlobalEnv>
```

##### KDF0

```
# Compute barcode rank
bc_rank <- barcodeRanks(KDF0)

# make knee plot
qplot(bc_rank$total, bc_rank$rank, geom = "line") +
  scale_x_log10() +
  scale_y_log10() +
  labs(y = "Total UMI count", x = "Barcode rank")
```

```
## Warning: `qplot()` was deprecated in ggplot2 3.4.0.
## This warning is displayed once every 8 hours.
## Call `lifecycle::last_lifecycle_warnings()` to see where this warning was
## generated.
```

```
## Warning in scale_x_log10(): log-10 transformation introduced infinite values.
```

###### Perform QC and filter

```
KDF0 <- subset(KDF0, subset =  nCount_RNA > 500)
#make scatter plots
plot1 <- FeatureScatter(KDF0, feature1 = "nCount_RNA", feature2 = "percent.mt")
```

```
## Warning: The `slot` argument of `FetchData()` is deprecated as of SeuratObject 5.0.0.
## ℹ Please use the `layer` argument instead.
## ℹ The deprecated feature was likely used in the Seurat package.
##   Please report the issue at <https://github.com/satijalab/seurat/issues>.
## This warning is displayed once every 8 hours.
## Call `lifecycle::last_lifecycle_warnings()` to see where this warning was
## generated.
```

```
plot2 <- FeatureScatter(KDF0, feature1 = "nCount_RNA", feature2 = "nFeature_RNA")
plot1 + plot2
```

```
#filtering
KDF0 <- subset(KDF0, subset =  percent.mt < 30 & nCount_RNA < 20000 & nFeature_RNA > 500 & nFeature_RNA < 5000)


VlnPlot(KDF0, features = c("nFeature_RNA", "nCount_RNA", "percent.mt"), ncol = 3)
```

```
## Warning: `PackageCheck()` was deprecated in SeuratObject 5.0.0.
## ℹ Please use `rlang::check_installed()` instead.
## ℹ The deprecated feature was likely used in the Seurat package.
##   Please report the issue at <https://github.com/satijalab/seurat/issues>.
## This warning is displayed once every 8 hours.
## Call `lifecycle::last_lifecycle_warnings()` to see where this warning was
## generated.
```

```
#check how many cells left
dim(KDF0)
```

```
## [1] 32443  9865
```

##### KDF4

```
# Compute barcode rank
bc_rank <- barcodeRanks(KDF4)

# make knee plot
qplot(bc_rank$total, bc_rank$rank, geom = "line") +
  scale_x_log10() +
  scale_y_log10() +
  labs(y = "Total UMI count", x = "Barcode rank")
```

```
## Warning in scale_x_log10(): log-10 transformation introduced infinite values.
```

###### Perform QC and filter

```
KDF4 <- subset(KDF4, subset =  nCount_RNA > 500)

#make scatter plots
plot1 <- FeatureScatter(KDF4, feature1 = "nCount_RNA", feature2 = "percent.mt")
plot2 <- FeatureScatter(KDF4, feature1 = "nCount_RNA", feature2 = "nFeature_RNA")
plot1 + plot2
```

```
#filtering
KDF4 <- subset(KDF4, subset =  percent.mt < 6 & nCount_RNA > 2000 & nCount_RNA < 30000 & nFeature_RNA > 2000 & nFeature_RNA < 6000)

# use Violin plots to adjust cut-offs as needed (remove outliers and low Features/counts)
VlnPlot(KDF4, features = c("nFeature_RNA", "nCount_RNA", "percent.mt"), ncol = 3)
```

```
#check how many cells left
dim(KDF4)
```

```
## [1] 29720  7041
```

##### NDF0

```
# Compute barcode rank
bc_rank <- barcodeRanks(NDF0)

# make knee plot
qplot(bc_rank$total, bc_rank$rank, geom = "line") +
  scale_x_log10() +
  scale_y_log10() +
  labs(y = "Total UMI count", x = "Barcode rank")
```

```
## Warning in scale_x_log10(): log-10 transformation introduced infinite values.
```

###### Perform QC and filter

```
NDF0 <- subset(NDF0, subset =  nCount_RNA > 500)

#make scatter plots
plot1 <- FeatureScatter(NDF0, feature1 = "nCount_RNA", feature2 = "percent.mt")
plot2 <- FeatureScatter(NDF0, feature1 = "nCount_RNA", feature2 = "nFeature_RNA")
plot1 + plot2
```

```
#filtering
NDF0 <- subset(NDF0, subset = percent.mt > 1 & percent.mt < 30 & nCount_RNA < 20000 & nFeature_RNA < 5000 & nFeature_RNA > 500)

# use Violin plots to adjust cut-offs as needed (remove outliers and low Features/counts)
VlnPlot(NDF0, features = c("nFeature_RNA", "nCount_RNA", "percent.mt"), ncol = 3)
```

```
#check how many cells left
dim(NDF0)
```

```
## [1] 32699  9845
```

##### NDF4

```
# Compute barcode rank
bc_rank <- barcodeRanks(NDF4)

# make knee plot
qplot(bc_rank$total, bc_rank$rank, geom = "line") +
  scale_x_log10() +
  scale_y_log10() +
  labs(y = "Total UMI count", x = "Barcode rank")
```

```
## Warning in scale_x_log10(): log-10 transformation introduced infinite values.
```

###### Perform QC and filter

```
NDF4 <- subset(NDF4, subset =  nCount_RNA > 500)

#make scatter plots
plot1 <- FeatureScatter(NDF4, feature1 = "nCount_RNA", feature2 = "percent.mt")
plot2 <- FeatureScatter(NDF4, feature1 = "nCount_RNA", feature2 = "nFeature_RNA")
plot1 + plot2
```

```
#filtering
NDF4 <- subset(NDF4, subset =   percent.mt < 7 & nCount_RNA > 2000 & nCount_RNA < 30000 & nFeature_RNA > 2000 & nFeature_RNA < 5500)

# use Violin plots to adjust cut-offs as needed (remove outliers and low Features/counts)
VlnPlot(NDF4, features = c("nFeature_RNA", "nCount_RNA", "percent.mt"), ncol = 3)
```

```
#check how many cells left
dim(NDF4)
```

```
## [1] 30032  8207
```

#### Merge all sample objects into one

```
## Merge all samples to your first sample object, making just one object of all samples
merged <- merge(x = KDF0, y = c(KDF4, NDF0, NDF4), add.cell.ids = c("KDF0", "KDF4", "NDF0", "NDF4"), project = "merged")

#save filtered, non-normalised seurat object
#save(merged, file= "processed.Robj")
load(file= "processed.Robj")
```

#### Run normalisation and scTransform with cell cycle regression by satija lab:

https://satijalab.org/seurat/

```
# clear environment to help with processing
rm(KDF0,KDF4, NDF0, NDF4, plot1, plot2, res_mat, seu, seurat_list, t2g, ens2sym, merged)  # Remove specific Seurat objects
gc()  # Force garbage collection

## Read in canonical cell cycle markers
s.genes <- cc.genes$s.genes
g2m.genes <- cc.genes$g2m.genes

sct_v4 <- NormalizeData(merged_tight)
```

```
## Warning: The `slot` argument of `SetAssayData()` is deprecated as of SeuratObject 5.0.0.
## ℹ Please use the `layer` argument instead.
## ℹ The deprecated feature was likely used in the Seurat package.
##   Please report the issue at <https://github.com/satijalab/seurat/issues>.
## This warning is displayed once every 8 hours.
## Call `lifecycle::last_lifecycle_warnings()` to see where this warning was
## generated.
```

```
sct_v4 <- CellCycleScoring(sct_v4,s.features = s.genes,g2m.features = g2m.genes)
```

```
## Warning: The following features are not present in the object: MLF1IP, not
## searching for symbol synonyms
```

```
## Warning: The following features are not present in the object: FAM64A, HN1, not
## searching for symbol synonyms
```

```
sct_v4 <- SCTransform(sct_v4, vst.flavor = "V2", vars.to.regress = c("S.Score", "G2M.Score"), verbose = TRUE)
```

```
## Calculating cell attributes from input UMI matrix: log_umi
```

```
## Variance stabilizing transformation of count matrix of size 33107 by 34948
```

```
## Model formula is y ~ log_umi
```

```
## Get Negative Binomial regression parameters per gene
```

```
## Using 2000 genes, 5000 cells
```

```
## Found 105 outliers - those will be ignored in fitting/regularization step
```

```
## Second step: Get residuals using fitted parameters for 33107 genes
```

```
## Computing corrected count matrix for 33107 genes
```

```
## Calculating gene attributes
```

```
## Wall clock passed: Time difference of 8.059657 mins
```

```
## Determine variable features
```

```
## Place corrected count matrix in counts slot
```

```
## Regressing out S.Score, G2M.Score
```

```
## Centering data matrix
```

```
## Set default assay to SCT
```

#### Run PCA and Elbow plot to determine PCs to take forward

```
sct_v4 <- RunPCA(sct_v4, npcs = 30)
```

```
## PC_ 1 
## Positive:  HSPA1A, CD74, HLA-DRB1, DNAJB1, HLA-DRA, SRGN, CD69, GPR183, NFKBIA, HLA-DPA1 
##     JUNB, HLA-DPB1, HSPA1B, DUSP2, MT-RNR1, HLA-DQB1, TNFAIP3, IL32, HSP90AA1, CD52 
##     BTG1, JUND, LTB, CXCR4, RGS1, NR4A2, ZFP36, CCL5, PTPRC, HLA-DQA1 
## Negative:  LGALS1, VIM, S100A6, COL1A1, COL1A2, IGFBP3, FN1, GAPDH, TAGLN, TPM2 
##     ANXA2, TMSB10, FTL, MT2A, TPM1, LDHA, THBS1, S100A4, SERPINE2, COL3A1 
##     S100A11, IGFBP6, GREM1, CALD1, C12orf75, TNFRSF11B, DCN, MMP3, CD63, DKK1 
## PC_ 2 
## Positive:  LGALS1, VIM, IGFBP3, S100A6, TMSB10, GAPDH, FN1, FTL, ANXA2, TMSB4X 
##     SH3BGRL3, GREM1, CD69, SERPINE2, TNFRSF11B, MMP3, LDHA, FTH1, DKK1, IGFBP6 
##     C12orf75, THBS1, SRGN, GPR183, S100A4, DUSP2, LOX, S100A11, CEMIP, PFN1 
## Negative:  MGP, APOE, IGFBP7, RGS5, ACTA2, SPARCL1, COL3A1, C11orf96, MT-RNR1, NDUFA4L2 
##     COL4A1, A2M, IGFBP5, RGS16, DCN, COL14A1, COL4A2, COL18A1, CFD, ID4 
##     C1R, SERPINF1, CPE, CFH, SOD3, MYH11, SFRP2, NR2F2, ABCC9, CXCL14 
## PC_ 3 
## Positive:  CD69, IL32, DNAJB1, HSPA1A, LTB, CD52, CCL5, HSP90AA1, IL7R, GZMA 
##     JUND, TRBC1, TRBC2, KLRB1, CD3D, BTG1, DUSP2, TRAC, PTPRC, CD2 
##     RGCC, HSPA1B, TNFAIP3, CXCR4, CD3E, TSC22D3, JUNB, CORO1A, CD3G, IFNG 
## Negative:  CD74, HLA-DRA, HLA-DRB1, HLA-DPA1, HLA-DPB1, CLDN5, AQP1, PECAM1, ACKR1, SPARCL1 
##     HLA-DQB1, HLA-DQA1, LYZ, PLVAP, AIF1, CXCL8, HLA-DMA, TYROBP, CD34, VWF 
##     RNASE1, SOX18, IL1B, RAMP2, SELE, CST3, FCER1A, EGFL7, IFI27, EMCN 
## PC_ 4 
## Positive:  DCN, SFRP2, CFD, COL3A1, PTGDS, COL1A1, COL1A2, CXCL14, LUM, FBLN1 
##     OGN, POSTN, C1R, CTSK, APOE, CCN5, ASPN, APOD, C1S, MFAP4 
##     SERPINF1, SPARC, C3, CTHRC1, CST3, COL6A2, COMP, FGL2, LYZ, TYROBP 
## Negative:  SPARCL1, ACTA2, RGS5, CLDN5, PECAM1, AQP1, ACKR1, A2M, IGFBP7, PLVAP 
##     NDUFA4L2, VWF, TM4SF1, MYH11, TAGLN, SOX18, C11orf96, CD34, MCAM, ADIRF 
##     EGFL7, SELE, RAMP2, EMCN, ADGRL4, BCAM, SOCS3, RAMP3, IFI27, ECSCR 
## PC_ 5 
## Positive:  AQP1, CLDN5, PECAM1, ACKR1, DCN, SFRP2, PLVAP, PTGDS, CFD, CD34 
##     CXCL14, VWF, CLU, SOX18, RAMP2, SPARCL1, EGFL7, SELE, EMCN, ADGRL4 
##     IFI27, GIMAP7, RAMP3, TNFSF10, COL15A1, ECSCR, PALMD, LUM, CLEC14A, TM4SF1 
## Negative:  ACTA2, RGS5, LYZ, TYROBP, CXCL8, HLA-DQA1, AIF1, HLA-DPB1, TAGLN, NDUFA4L2 
##     HLA-DQB1, IL1B, HLA-DPA1, CD83, FCER1A, CCL3L1, FCER1G, GPR183, PLEK, CCL3 
##     C11orf96, CLEC10A, MYH11, IGFBP7, MS4A6A, BCL2A1, MNDA, FCGR2A, COL4A1, LGALS2
```

```
ElbowPlot(sct_v4, ndims = 30)
```

#### Run unsupervised clustering

```
sct_v4 <- FindNeighbors(sct_v4, dims = 1:20, reduction = "pca")
```

```
## Computing nearest neighbor graph
```

```
## Computing SNN
```

```
sct_v4 <- FindClusters(sct_v4, resolution = 0.2)
#11 clusters
table(sct_v4$orig.ident)
sct_v4 <- RunUMAP(sct_v4, reduction = "pca",  dims = 1:20)
```

```
## Warning: The default method for RunUMAP has changed from calling Python UMAP via reticulate to the R-native UWOT using the cosine metric
## To use Python UMAP via reticulate, set umap.method to 'umap-learn' and metric to 'correlation'
## This message will be shown once per session
```

```
## 13:43:15 UMAP embedding parameters a = 0.9922 b = 1.112
```

```
## 13:43:15 Read 34948 rows and found 20 numeric columns
```

```
## 13:43:15 Using Annoy for neighbor search, n_neighbors = 30
```

```
## 13:43:15 Building Annoy index with metric = cosine, n_trees = 50
```

```
## 0%   10   20   30   40   50   60   70   80   90   100%
```

```
## [----|----|----|----|----|----|----|----|----|----|
```

```
## **************************************************|
## 13:43:17 Writing NN index file to temp file /var/folders/t_/9nwsb_7x4_q9zfch7sf3tv440000gn/T//RtmpkEpUBD/fileb526650b6913
## 13:43:17 Searching Annoy index using 1 thread, search_k = 3000
## 13:43:22 Annoy recall = 100%
## 13:43:24 Commencing smooth kNN distance calibration using 1 thread with target n_neighbors = 30
## 13:43:25 Initializing from normalized Laplacian + noise (using RSpectra)
## 13:43:26 Commencing optimization for 200 epochs, with 1455374 positive edges
## 13:43:26 Using rng type: pcg
## 13:43:33 Optimization finished
```

```
#save(sct_v4, file = "sct_v4.Robj")
load(file = "sct_v4.Robj")
```

#### Visualize UMAP

```
Idents(sct_v4) <- sct_v4$seurat_clusters
DimPlot(sct_v4, reduction = "umap", label=TRUE, split.by = "orig.ident")
```

#### Find marker genes and cell proportions for clusters

```
SCT_v4.markers <- FindAllMarkers(sct_v4, min.pct = 0.25,  only.pos=TRUE)
```

```
## Calculating cluster 0
```

```
## Calculating cluster 1
```

```
## Calculating cluster 2
```

```
## Calculating cluster 3
```

```
## Calculating cluster 4
```

```
## Calculating cluster 5
```

```
## Calculating cluster 6
```

```
## Calculating cluster 7
```

```
## Calculating cluster 8
```

```
## Calculating cluster 9
```

```
## Calculating cluster 10
```

```
write.table(SCT_v4.markers, "SCT_v4.markers_unintegrated.txt")

props <- table(sct_v4$seurat_clusters, sct_v4$orig.ident)
write.table(props, file = 'V4_SCT_nointeg_props.txt')
```

#### Renaming clusters as cell types

```
Idents(sct_v4) <- sct_v4$seurat_clusters
levels(sct_v4)
paper_clusters <- c("c0 Fibroblast", "c1 SMC/Myofibroblast", "c2 T-cell", "c0 Fibroblast", "c0 Fibroblast", "c3 NK cell", "c4 Endothelial cell", "c5 Dendritic cell", "c6 B cell", "c7 Mast cell", "c8 Schwann cell")

names(paper_clusters) <- levels(sct_v4)
sct_v4 <- RenameIdents(sct_v4, paper_clusters)
sct_v4[["paper_clusters"]] <- Idents(object = sct_v4) #to add clusters to metadata.
```

#### Creating cell-type UMAP split by sample

```
#defining cluster colours
cluster_colors <- c("c0 Fibroblast" = "#99CC33", "c4 Endothelial cell" = "#FF3399", "c3 NK cell" = "orange", "c1 SMC/Myofibroblast" = "darkturquoise", "c2 T-cell" = "red", "c5 Dendritic cell"="#00ccff", "c8 Schwann cell"="pink", "c7 Mast cell"="aquamarine" , "c6 B cell"="purple") 

DimPlot(sct_v4, reduction = "umap", split.by = "orig.ident", label = FALSE)+theme(legend.text = element_text(size = 20), axis.text = element_text(size = 40, colour = "black"), axis.title = element_text(size = 50))+ scale_color_manual(values = cluster_colors)
```

#### Marker Dotplot

```
### DEGs markers for each cluster

DotPlot(sct_v4, features = c("THY1", "CEMIP", "RGS5", "ACTA2", "PTPRC", "CD3D","TRDC","IL32","PECAM1", "ACKR1","LYZ", "IL1B", "CD79A", "IGHM", "TPSB2", "KIT", "SOX10", "S100B"), dot.scale = 20) +theme(text = element_text(size = 25, face = "bold"), axis.text = element_text(size = 40, colour = "black"), axis.title = element_text(size = 50)) + RotatedAxis()
```

#### Get proportions per cell type

```
proportions <- table(sct_v4$paper_clusters, sct_v4$orig.ident)
```

#### Stromal subsetting

```
Idents(sct_v4) <- sct_v4$paper_clusters
sct_v4_stromal <- subset(x = sct_v4, idents = (c("c0 Fibroblast","c1 SMC/Myofibroblast", "c4 Endothelial cell")))
DefaultAssay(sct_v4_stromal) <- "SCT"

sct_v4_stromal <- RunPCA(sct_v4_stromal, npcs = 30)
```

```
## PC_ 1 
## Positive:  SPARCL1, MGP, APOE, RGS5, MT-RNR1, IGFBP7, ACTA2, C11orf96, A2M, HSPA1A 
##     NDUFA4L2, COL4A1, RGS16, COL3A1, MT-RNR2, SOCS3, FOS, AQP1, JUNB, JUN 
##     COL4A2, ZFP36, IGFBP5, COL18A1, CLDN5, ID4, MYH11, MT-ND3, CD74, CFH 
## Negative:  LGALS1, IGFBP3, VIM, S100A6, FN1, GAPDH, TMSB10, FTL, ANXA2, GREM1 
##     SERPINE2, THBS1, TNFRSF11B, IGFBP6, MMP3, C12orf75, DKK1, LDHA, LOX, TPM2 
##     S100A11, SH3BGRL3, S100A4, SERPINE1, ADM, CEMIP, PKM, MT2A, FTH1, POLR2L 
## PC_ 2 
## Positive:  CLDN5, PECAM1, ACKR1, AQP1, CD74, HLA-DRB1, SPARCL1, PLVAP, HLA-DRA, VWF 
##     CD34, SOX18, SELE, RAMP2, EGFL7, RNASE1, EMCN, ADGRL4, TM4SF1, RAMP3 
##     GIMAP7, ECSCR, MCTP1, PALMD, C2CD4B, CD93, CLEC14A, IFI27, CLU, SOCS3 
## Negative:  DCN, COL3A1, SFRP2, COL1A1, COL1A2, APOE, CFD, PTGDS, CXCL14, MGP 
##     LUM, C1R, SERPINF1, SPARC, POSTN, COL14A1, FBLN1, ASPN, COL6A2, C1S 
##     CCN5, OGN, CTSK, MFAP4, COL6A1, APOD, COL5A2, C3, COMP, RARRES2 
## PC_ 3 
## Positive:  DCN, SFRP2, CFD, PTGDS, CXCL14, COL1A2, COL1A1, COL3A1, LUM, AQP1 
##     CLDN5, ACKR1, PECAM1, FBLN1, CD74, OGN, HLA-DRB1, APOD, POSTN, CD34 
##     CTSK, CLU, PLVAP, CTHRC1, HLA-DRA, CCN5, VWF, C3, ASPN, RAMP2 
## Negative:  ACTA2, RGS5, NDUFA4L2, TAGLN, MYH11, C11orf96, IGFBP7, COL4A1, RGS16, MCAM 
##     ID4, COL4A2, NOTCH3, NR2F2, HOPX, COL18A1, PPP1R14A, STEAP4, SYNPO2, ABCC9 
##     CPE, CALD1, TINAGL1, ADIRF, COX4I2, RERGL, MYL9, HIGD1B, EGFL6, FMO3 
## PC_ 4 
## Positive:  HSPA1A, CD69, DNAJB1, IL32, LTB, CFD, CD52, SRGN, GZMA, DUSP2 
##     CXCL14, CCL5, TNFAIP3, GPR183, HSP90AA1, PTPRC, MT-RNR1, JUND, JUNB, CXCR4 
##     HSPA1B, TRBC1, BTG1, NFKBIA, KLRB1, TRBC2, RGS1, HCST, CORO1A, APOD 
## Negative:  COL1A1, SPARCL1, ACTA2, COL1A2, COL3A1, SPARC, IGFBP7, COL4A1, TAGLN, POSTN 
##     AQP1, CLDN5, A2M, PECAM1, ACKR1, FN1, COL4A2, IGFBP3, CTHRC1, PLVAP 
##     IFI27, CALD1, COL14A1, MGP, RGS5, TPM2, LGALS1, VIM, VWF, ASPN 
## PC_ 5 
## Positive:  COL1A1, COL3A1, COL1A2, SPARC, POSTN, ASPN, CTHRC1, LUM, COL5A2, SFRP4 
##     COMP, MDK, HSPA1A, COL14A1, COL6A1, CD69, COL5A1, COL11A1, COL4A1, LTB 
##     DNAJB1, COL6A2, IL32, C1QTNF3, SRGN, CD52, GZMA, CPXM1, DUSP2, TNFAIP3 
## Negative:  CFD, CXCL14, APOD, APOE, C3, PTGDS, IGFBP5, ABCA8, CXCL12, CFH 
##     GSN, SELENOP, SOD3, ADH1B, EFEMP1, DCN, VIM, CCL19, PLA2G2A, C11orf96 
##     FGL2, C1R, PI16, ABCA10, C7, ADIRF, CCDC80, SPARCL1, C1S, TNXB
```

```
ElbowPlot(sct_v4_stromal, ndims = 30)
```

```
sct_v4_stromal <- FindNeighbors(sct_v4_stromal, dims = 1:20, reduction = "pca")
```

```
## Computing nearest neighbor graph
```

```
## Computing SNN
```

```
sct_v4_stromal <- FindClusters(sct_v4_stromal, resolution = 0.30)
#8 clusters
sct_v4_stromal <- RunUMAP(sct_v4_stromal, reduction = "pca",  dims = 1:20)
```

```
## 13:52:12 UMAP embedding parameters a = 0.9922 b = 1.112
```

```
## 13:52:12 Read 25956 rows and found 20 numeric columns
```

```
## 13:52:12 Using Annoy for neighbor search, n_neighbors = 30
```

```
## 13:52:12 Building Annoy index with metric = cosine, n_trees = 50
```

```
## 0%   10   20   30   40   50   60   70   80   90   100%
```

```
## [----|----|----|----|----|----|----|----|----|----|
```

```
## **************************************************|
## 13:52:13 Writing NN index file to temp file /var/folders/t_/9nwsb_7x4_q9zfch7sf3tv440000gn/T//RtmpkEpUBD/fileb526505a1e48
## 13:52:13 Searching Annoy index using 1 thread, search_k = 3000
## 13:52:17 Annoy recall = 100%
## 13:52:18 Commencing smooth kNN distance calibration using 1 thread with target n_neighbors = 30
## 13:52:19 Initializing from normalized Laplacian + noise (using RSpectra)
## 13:52:19 Commencing optimization for 200 epochs, with 1060366 positive edges
## 13:52:19 Using rng type: pcg
## 13:52:25 Optimization finished
```

```
#save(sct_v4_stromal, file = "sct_v4_stromal.Robj")
load(file = "sct_v4_stromal.Robj")
```

##### Visualise UMAP

```
Idents(sct_v4_stromal) <- sct_v4_stromal$seurat_clusters
DimPlot(sct_v4_stromal, reduction = "umap", split.by = "orig.ident")
```

##### Find markers and rename clusters

```
sct_v4_stromal.markers <- FindAllMarkers(sct_v4_stromal, min.pct = 0.25,  only.pos=TRUE)
```

```
## Calculating cluster 0
```

```
## Calculating cluster 1
```

```
## Calculating cluster 2
```

```
## Calculating cluster 3
```

```
## Calculating cluster 4
```

```
## Calculating cluster 5
```

```
## Calculating cluster 6
```

```
## Calculating cluster 7
```

```
write.table(sct_v4_stromal.markers, "sct_v4_stromal.markers_unintegrated.txt")

Idents(sct_v4_stromal) <- sct_v4_stromal$seurat_clusters
levels(sct_v4_stromal)
paper_clusters <- c("c0 Fibroblast CEMIP+", "c1 SMC/Myofibroblast", "c2 Proliferating fibroblast", "c2 Proliferating fibroblast", "c1 SMC/Myofibroblast", "c3 Endothelial cell", "c4 Fibroblast FAP+/SFRP2+", "c5 Fibroblast SFRP1+")


names(paper_clusters) <- levels(sct_v4_stromal)
sct_v4_stromal <- RenameIdents(sct_v4_stromal, paper_clusters)
sct_v4_stromal[["paper_clusters"]] <- Idents(object = sct_v4_stromal) #to add clusters to metadata. 
Idents(sct_v4_stromal) <- sct_v4_stromal$paper_clusters
```

##### Make colour-coded UMAP

```
#defining cluster colours
cluster_colors <- c("c0 Fibroblast CEMIP+" = "#99FF33","c3 Endothelial cell" = "#FF3399", "c4 Fibroblast FAP+/SFRP2+" = "#99CC33", "c1 SMC/Myofibroblast" = "darkturquoise", "c5 Fibroblast SFRP1+" = "#009933", "c2 Proliferating fibroblast"="darkblue")  # customize as needed

DimPlot(sct_v4_stromal, reduction = "umap", split.by = "orig.ident", label = FALSE)+theme(legend.text = element_text(size = 20), axis.text = element_text(size = 40, colour = "black"), axis.title = element_text(size = 50))+ scale_color_manual(values = cluster_colors)
```

##### Marker Dotplot

```
### DEGs markers for each cluster

DotPlot(sct_v4_stromal, features = c("THY1","CEMIP","LMCD1","RGS5", "NOTCH3","MCAM","ACTA2", "CENPF", "FABP5", "PECAM1", "ACKR1", "FAP","SFRP2", "SFRP1","OGN", "PI16","CXCL10", "SPARC","COL3A1","LRRC15"), dot.scale = 20)+theme(text = element_text(size = 25, face = "bold"), axis.text = element_text(size = 40, colour = "black"), axis.title = element_text(size = 50)) + RotatedAxis()
```

```
# Markers for Sole-Boldo et al. Buechler et al. and Korsunsky et al. 
DotPlot(sct_v4_stromal, features = c("THY1", "CCN5","SLPI","CCL19", "APOE", "CXCL2","APCDD1", "WIF1","PTGDS", "ASPN", "POSTN","COMP","GPC3","TNN", "SFRP1", "PI16","CXCL10", "SPARC","COL3A1","LRRC15"), dot.scale = 20, col.min = 0.5)+theme(text = element_text(size = 30, face = "bold"), axis.text = element_text(size = 30, colour = "black"), axis.title = element_text(size = 50)) + RotatedAxis()
```

##### Making stacked barcharts of cell cluster proportions, by sample

```
Idents(sct_v4_stromal) <- sct_v4_stromal$orig.ident
levels(sct_v4_stromal)
```

```
## [1] "KDF0" "KDF4" "NDF0" "NDF4"
```

```
AL_ident <- c("K.P0", "K.P4", "N.P0", "N.P4")

names(AL_ident) <- levels(sct_v4_stromal)
sct_v4_stromal <- RenameIdents(sct_v4_stromal, AL_ident)
sct_v4_stromal[["AL_ident"]] <- Idents(object = sct_v4_stromal) #to add clusters to metadata. 


Idents(sct_v4_stromal) <- sct_v4_stromal$paper_clusters
pt <- prop.table(table(Idents(sct_v4_stromal), sct_v4_stromal$AL_ident), margin = 2)
pt <- as.data.frame(pt)

colnames(pt)
```

```
## [1] "Var1" "Var2" "Freq"
```

```
plot <- ggplot(pt, aes(x = Var2, y = Freq, fill = Var1)) +
  theme_bw(base_size = 15) +
  geom_col(position = "fill", width = 0.8) +
  xlab("Sample") +
  ylab("Proportion") +
  scale_fill_manual(values = brewer.pal(12, "Paired")) +
  theme(legend.title = element_blank())


# plot
plot
```

```
#plot + theme(text = element_text(size = 50, face = "bold"), axis.text = element_text(size = 30, colour = "black"), axis.text.x = element_text(angle = 0, vjust = 0.5, hjust=1))
```

#### P4-only subclustering

```
P4 <- subset(x = sct_v4_stromal, idents = (c("c0 Fibroblast CEMIP+", "c2 Proliferating fibroblast")))
Idents(P4) <- P4$orig.ident
P4 <- subset(x = P4, idents = (c("KDF4", "NDF4")))
DefaultAssay(P4) <- "SCT"

P4 <- RunPCA(P4, npcs = 50)
```

```
## PC_ 1 
## Positive:  MT-RNR1, HSPA1A, MGP, SPARCL1, APOE, RGS5, IGFBP7, ACTA2, JUNB, DNAJB1 
##     HSPA1B, MT-RNR2, JUN, FOS, CD74, C11orf96, CD69, ZFP36, NFKBIA, HLA-DRB1 
##     HLA-B, COL3A1, MT-ND3, B2M, IL32, A2M, MT-ATP6, HLA-DRA, NDUFA4L2, DUSP1 
## Negative:  LGALS1, IGFBP3, VIM, S100A6, FN1, GAPDH, TMSB10, ANXA2, THBS1, FTL 
##     SERPINE2, GREM1, TPM2, TNFRSF11B, IGFBP6, MMP3, LDHA, C12orf75, DKK1, LOX 
##     S100A11, MT2A, SERPINE1, S100A4, TPM1, ADM, CEMIP, PKM, SH3BGRL3, POLR2L 
## PC_ 2 
## Positive:  MT2A, VIM, TUBA1B, TMSB10, C12orf75, MT1E, STMN1, LGALS1, PTTG1, S100A10 
##     JPT1, LMNA, CAV1, CDKN3, HMGN2, DRAP1, ANXA2, CLDN11, TUBB4B, PPP1R14B 
##     S100A6, H2AFZ, DKK1, CXCL12, SMS, RPL22L1, BIRC5, GRP, EIF5A, TK1 
## Negative:  IGFBP3, FN1, FTL, THBS1, FTH1, TNFRSF11B, TIMP3, COL1A1, ELN, COL1A2 
##     GREM1, COL12A1, MMP3, ACTC1, STC1, TAGLN, POSTN, CALD1, COMP, TPM1 
##     CYP1B1, KRT7, SULF1, CCN2, TPM2, IGFBP5, LMCD1, HIST1H4C, NUPR1, ACAN 
## PC_ 3 
## Positive:  SERPINE2, PTX3, MFAP5, FN1, SERPINE1, HAS2, KRT19, ACAN, CCDC80, CEMIP 
##     COL1A1, HIST1H4C, HAPLN1, THBS1, INHBA, FBN2, CALD1, TAGLN, IGFBP5, GREM1 
##     AKAP12, TOP2A, OXTR, LOX, ITGB1, COL8A1, COL11A1, TPM2, CTHRC1, CLDN11 
## Negative:  MMP3, IGFBP3, FTL, FTH1, ACTC1, LY6K, DKK1, NQO1, PSG5, KRT7 
##     AKR1C1, TXN, STMN2, PTTG1, ADM, GAPDH, S100A4, CLEC2B, CLIC3, LINC01615 
##     EMX2OS, CRYAB, LGALS1, GZMA, CDKN3, OSR2, WFDC21P, STMN1, CKB, SPON2 
## PC_ 4 
## Positive:  MMP3, ACTC1, DKK1, VIM, STMN2, THBS1, KRT7, C12orf75, CLIC3, CRYAB 
##     TNFRSF11B, APLP2, DAB2, NEAT1, CXCL12, PSG5, TGFBI, S100A4, DST, TIMP3 
##     F3, TPM1, LMO7, KRT19, NEK7, TENM2, SSX2IP, GRP, TOP2A, OXTR 
## Negative:  FTL, FTH1, SERPINE2, MT2A, POSTN, GAPDH, MFAP5, NUPR1, COL1A1, MGST1 
##     S100A6, FN1, NQO1, PSAT1, TXN, PHGDH, AKR1C1, LINC01615, TIMP1, EIF4EBP1 
##     RPS18, EEF1A1, RPL41, CTHRC1, RPS19, RPS12, TPI1, GAS6, HMOX1, RPS27 
## PC_ 5 
## Positive:  IGFBP3, KRT19, POSTN, SERPINE1, KRTAP2-3, CCDC80, LOX, PSG5, TPM1, LOXL2 
##     COL1A1, FN1, ENPP2, TENM2, BNIP3, CLDN11, MT2A, KRT34, COL1A2, LY6K 
##     NRG1, GLIPR1, PTX3, HSP90B1, TMSB10, HSPA5, SMS, HIF1A, ENO1, DKK1 
## Negative:  ACAN, MMP3, STMN2, MFAP5, TNFRSF11B, CYP1B1, IGFBP2, SERPINE2, PPP1R14A, SULF1 
##     S100A4, COMP, FBXO32, HAPLN1, IGFBP6, TIMP1, COL12A1, FTH1, GRP, PRELP 
##     DCN, LGALS1, LUM, MFGE8, COL11A1, FTL, IGFBP4, S100A6, MME, APLP2
```

```
ElbowPlot(P4, ndims = 30)
```

```
P4 <- FindNeighbors(P4, dims = 1:20, reduction = "pca")
```

```
## Computing nearest neighbor graph
```

```
## Computing SNN
```

```
P4 <- FindClusters(P4, resolution = 0.15)
#5 clusters
P4 <- RunUMAP(P4, reduction = "pca",  dims = 1:20)
```

```
## 13:54:56 UMAP embedding parameters a = 0.9922 b = 1.112
```

```
## 13:54:56 Read 15240 rows and found 20 numeric columns
```

```
## 13:54:56 Using Annoy for neighbor search, n_neighbors = 30
```

```
## 13:54:56 Building Annoy index with metric = cosine, n_trees = 50
```

```
## 0%   10   20   30   40   50   60   70   80   90   100%
```

```
## [----|----|----|----|----|----|----|----|----|----|
```

```
## **************************************************|
## 13:54:57 Writing NN index file to temp file /var/folders/t_/9nwsb_7x4_q9zfch7sf3tv440000gn/T//RtmpkEpUBD/fileb5265a56ad92
## 13:54:57 Searching Annoy index using 1 thread, search_k = 3000
## 13:54:59 Annoy recall = 100%
## 13:55:00 Commencing smooth kNN distance calibration using 1 thread with target n_neighbors = 30
## 13:55:01 Initializing from normalized Laplacian + noise (using RSpectra)
## 13:55:01 Commencing optimization for 200 epochs, with 649210 positive edges
## 13:55:01 Using rng type: pcg
## 13:55:04 Optimization finished
```

```
#save(P4, file = "P4_only.Robj")
load(file = "P4_only.Robj")
```

##### Visualise UMAP

```
Idents(P4) <- P4$seurat_clusters
DimPlot(P4, reduction = "umap", split.by = "orig.ident")
```

```
# 5 clusters
```

##### Find markers and rename clusters

```
P4.markers <- FindAllMarkers(P4, min.pct = 0.25,  only.pos=TRUE)
```

```
## Calculating cluster 0
```

```
## Calculating cluster 1
```

```
## Calculating cluster 2
```

```
## Calculating cluster 3
```

```
## Calculating cluster 4
```

```
write.table(P4.markers, "sct_v4_P4.markers.txt")

Idents(P4) <- P4$seurat_clusters
paper_clusters <- c("c0 CEMIP+", "c1 Proliferating", "c1 Proliferating", "c2 KRT19+", "c3 SRGN+")

names(paper_clusters) <- levels(P4)
P4 <- RenameIdents(P4, paper_clusters)
P4[["paper_clusters"]] <- Idents(object = P4) #to add clusters to metadata.
```

##### Marker Dotplot

```
DotPlot(P4 , features = c("CEMIP","CEBPD", "FGF7", "S.Score", "G2M.Score", 'CDC20','HIST1H1B','CDK1', "KRT19", "SRGN", "SVIL", "NTN4","PDGFRA", "DKK1","EPCAM"), dot.scale = 20)+theme(text = element_text(size = 25, face = "bold"), axis.text = element_text(size = 40, colour = "black"), axis.title = element_text(size = 50)) + RotatedAxis()
```

```
## Warning: Scaling data with a low number of groups may produce misleading
## results
```

##### Making stacked barcharts of cell cluster proportions, by sample

```
Idents(P4) <- P4$orig.ident
AL_ident <- c("K.P4", "N.P4")

names(AL_ident) <- levels(P4)
P4 <- RenameIdents(P4, AL_ident)
P4[["AL_ident"]] <- Idents(object = P4) #to add clusters to metadata. 


Idents(P4) <- P4$paper_clusters
pt <- prop.table(table(Idents(P4), P4$AL_ident), margin = 2)
pt <- as.data.frame(pt)

colnames(pt)
```

```
## [1] "Var1" "Var2" "Freq"
```

```
plot <- ggplot(pt, aes(x = Var2, y = Freq, fill = Var1)) +
  theme_bw(base_size = 15) +
  geom_col(position = "fill", width = 0.8) +
  xlab("Sample") +
  ylab("Proportion") +
  scale_fill_manual(values = brewer.pal(12, "Paired")) +
  theme(legend.title = element_blank())


# plot
plot
```

```
#plot + theme(text = element_text(size = 50, face = "bold"), axis.text = element_text(size = 40, colour = "black"), axis.text.x = element_text(angle = 0, vjust = 0.5, hjust=1))
```

##### Making gene signatures for P4 clusters and adding signature scores to stromal dataset

```
library(scCustomize)


#CEMIP + KRT19 >40% >1.5 ratio
#SRGN >60%, >3 ratio
#prolif >70%, >3 ratio

P4_signatures <- list(
CEMIPpos = c(
  "SYNPO2", "LMCD1", "NR2F1", "CEBPD", "ITGA11", "PNRC1", "VSIR", "THBS2",
  "LINC01615", "YPEL3", "HSPB7", "ZNF503", "STC2", "GLRX", "CEMIP", "MIR210HG",
  "ALPK2", "ZMAT3", "SLFN5", "LBH", "PLPP3", "FGF7", "RABGAP1", "FCGRT",
  "TMEM119", "GJA1", "C1orf198", "MXD4", "HERPUD1", "GREM2", "CDC42EP5",
  "WNT5A", "CEBPB", "SULF1", "YPEL5", "GALNT5", "MAP1LC3A", "PDGFRB", "ARMCX3",
  "JUNB", "SNHG18", "PCDH18", "ID2"),
Proliferating = c(
"HIST1H1B", "CDK1", "AURKB", "HJURP", "FBXO5", "MXD3", "CKAP2L", "NCAPG",
  "RAD51AP1", "SPC25", "NDC80", "KIF11", "NUF2", "BARD1", "CCNA2", "UBE2C",
  "AURKA", "HMMR", "TACC3", "GTSE1", "CLSPN", "CENPE", "RACGAP1", "SGO2",
  "KNL1", "CDKN2D", "RRM2", "PBK", "KIF23", "TOP2A", "KIF20B", "CENPK",
  "FAM111A", "SHCBP1", "ASPM", "ATAD2", "ZWINT", "MKI67", "MAD2L1", "UBE2T",
  "CDC20", "CEP55", "ECT2", "PRC1", "MCM7", "CDCA3", "ANLN", "GMNN",
  "DNAJC9", "CKAP2", "TPX2", "CCNB1", "DHFR", "DLGAP5", "DIAPH3", "CENPM",
  "SPDL1", "CENPF"),
KRT19pos = c(
  'KRT19', "KRTAP2-3","KRTAP1-5","BAALC"),
SRGNpos = c(
  "MEST", "ABI3BP", "SRGN", "TM4SF1", "FOXC2", "MEOX2", "SVIL", "NTN4",
  "PPP1R14A", "CELF2", "F2R", "DSP", "CYP1B1", "RGS4", "ANKRD37", "BST1",
  "IFI27", "COL4A1", "INHBA", "SLC6A6", "CD9", "CAVIN2", "AKAP12"))


DefaultAssay(sct_v4_stromal) <-"RNA"

sct_v4_stromal <- AddModuleScore_UCell(sct_v4_stromal, features=P4_signatures, name=NULL)


FeaturePlot_scCustom(sct_v4_stromal, reduction = "umap", features = names(P4_signatures), min.cutoff = "q5", max.cutoff = "q95", order=TRUE, keep.scale="all", na_cutoff = 0.1)
```

```
## Warning: Some of the plotted features are from meta.data slot.
## • Please check that `na_cutoff` param is being set appropriately for those
##   features.
```

#### P0 only dataset

```
P0 <- subset(x = sct_v4_stromal, idents = (c("c1 SMC/Myofibroblast", "c3 Endothelial cell", "c4 Fibroblast FAP+/SFRP2+","c5 Fibroblast SFRP1+")))

levels(P0)
```

```
## [1] "c1 SMC/Myofibroblast"      "c3 Endothelial cell"      
## [3] "c4 Fibroblast FAP+/SFRP2+" "c5 Fibroblast SFRP1+"
```

##### Making gene signatures for P0 stromal clusters and plotting onto all stromal clusters

```
# all over 70% expression in specific cluster and ratio >2 compared to other clusters
#SFRP1: all over 50% expression in specific cluster and ratio >2 compared to all other clusters


signatures <- list(
  SMC_Myofibroblast = c(
  "NDUFA4L2", "RGS5", "SYNPO2", "HOPX", "ABCC9", "NOTCH3", "ACTA2", "TINAGL1", "ID4", "LBH", "MCAM", "MYLK","CPE", "TAGLN", "CRIP1", "CHCHD10", "TPM2", "PDGFRB", "NR2F2", "FILIP1L"),
Endothelial_cell = c(
  "ECSCR", "IL3RA", "RNASE1", "VWF", "MCTP1", "PLVAP", "GIMAP7", "ITGA6",
  "GIMAP4", "ADGRL4", "CLDN5", "RAMP3", "SOX18", "PECAM1", "ACKR1", "EGFL7",
  "CLEC14A", "HLA-DPA1", "EMCN", "HLA-DRB1", "HLA-DRA", "CD34", "RAMP2",
  "CD93", "JAM2", "CD74"),
Fibroblast_FAP_SFRP2 = c(
  "DPT", "MXRA5", "SFRP2", "CXCL14", "MRC2", "PTGDS", "FBLN1", "MMP2",
  "MEG3", "THBS2", "OLFML3", "CFD", "VCAN", "FBLN2", "MFAP4", "CTSK",
  "RCN3", "LRP1", "MXRA8", "RND3", "LUM", "CD248", "COL5A1", "GPNMB",
  "TMEM176B", "TMEM176A", "COPZ2", "CLEC11A", "DAB21", "CXCL12", "EFEMP2",
  "NUCB2", "FBN1", "FKBP10", "GAS6", "NBL1", "DCN", "HTRA1"),
Fibroblast_SFRP1 = c(
  "TNN", "DKK2", "SFRP1", "MFAP5", "EMID1", "OGN", "LIMCH", "PDGFRA",
  "SPON1", "CXCL14", "MXRA5", "CRABP2", "DPT", "FBLN5", "PTGDS",
  "FBLN1", "MMP2", "PRELP", "FGFR1", "MEG3", "CFD", "TRPS1", "MAFB",
  "LTBP4", "OLFML3", "MFAP4", "PTN", "COL15A2", "CDH11", "MEIS2",
  "COL16A1", "PRNP", "VCAN", "LINC00632", "LRP1", "SRPX", "CTSK",
  "SSPN", "LUM", "KLF4", "FBLN2", "PIK3R1", "MXRA8", "FAM3C"
))

DefaultAssay(sct_v4_stromal) <-"RNA"

sct_v4_stromal <- AddModuleScore_UCell(sct_v4_stromal, features=signatures, name=NULL)
```

```
## Warning: The following genes were not found and will be
##                         imputed to exp=0:
## * DAB21,LIMCH,COL15A2
```

```
head(sct_v4_stromal[[]])
```

```
##                      orig.ident nCount_RNA nFeature_RNA percent.mt      S.Score
## KF0_AAACCCAGTGATTCTG       KDF0       3582         1488   16.44333 -0.001308664
## KF0_AAACCCATCAGGGTAG       KDF0       3385         1374   22.48154 -0.059779647
## KF0_AAACCCATCCAAAGGG       KDF0       2706         1146   26.12712 -0.068580993
## KF0_AAACCCATCCGTATAG       KDF0       7548         2342   17.24960 -0.090403185
## KF0_AAACGAAAGAGATCGC       KDF0       1820          834   16.48352 -0.016890475
## KF0_AAACGAAAGCATTTGC       KDF0       8910         2773   23.97306 -0.025520149
##                        G2M.Score Phase nCount_SCT nFeature_SCT SCT_snn_res.0.2
## KF0_AAACCCAGTGATTCTG -0.06663831    G1       7492         1609               1
## KF0_AAACCCATCAGGGTAG -0.09580494    G1       7501         1556               1
## KF0_AAACCCATCCAAAGGG -0.13232253    G1       7258         1501               1
## KF0_AAACCCATCCGTATAG -0.16707876    G1       7951         2304               1
## KF0_AAACGAAAGAGATCGC -0.12724578    G1       7063         1414               1
## KF0_AAACGAAAGCATTTGC -0.12591636    G1       8537         2727               1
##                      seurat_clusters Endothelial_cell Fibroblast_FAP_SFRP2
## KF0_AAACCCAGTGATTCTG               4       0.02225641           0.07277193
## KF0_AAACCCATCAGGGTAG               1       0.00000000           0.03349123
## KF0_AAACCCATCCAAAGGG               1       0.00000000           0.12057018
## KF0_AAACCCATCCGTATAG               4       0.00000000           0.06085965
## KF0_AAACGAAAGAGATCGC               1       0.00000000           0.08761404
## KF0_AAACGAAAGCATTTGC               1       0.00000000           0.06108772
##                      Fibroblast_SFRP1 Proliferating  CEMIPpos KRT19pos
## KF0_AAACCCAGTGATTCTG       0.06511364     0.0000000 0.1280853        0
## KF0_AAACCCATCAGGGTAG       0.00000000     0.0000000 0.1154806        0
## KF0_AAACCCATCCAAAGGG       0.03793939     0.0000000 0.1286744        0
## KF0_AAACCCATCCGTATAG       0.00000000     0.0000000 0.1071085        0
## KF0_AAACGAAAGAGATCGC       0.03468939     0.0302931 0.1206279        0
## KF0_AAACGAAAGCATTTGC       0.03134848     0.0000000 0.1734961        0
##                         SRGNpos SMC_Myofibroblast       paper_clusters AL_ident
## KF0_AAACCCAGTGATTCTG 0.14695652         0.5262500 c1 SMC/Myofibroblast     K.P0
## KF0_AAACCCATCAGGGTAG 0.08184058         0.7425500 c1 SMC/Myofibroblast     K.P0
## KF0_AAACCCATCCAAAGGG 0.15907246         0.6890000 c1 SMC/Myofibroblast     K.P0
## KF0_AAACCCATCCGTATAG 0.04123188         0.7555833 c1 SMC/Myofibroblast     K.P0
## KF0_AAACGAAAGAGATCGC 0.10500000         0.3915500 c1 SMC/Myofibroblast     K.P0
## KF0_AAACGAAAGCATTTGC 0.13944928         0.8111500 c1 SMC/Myofibroblast     K.P0
```

```
FeaturePlot_scCustom(sct_v4_stromal, reduction = "umap", features = names(signatures), min.cutoff = "q10", max.cutoff = "q100", order=TRUE, keep.scale="all", na_cutoff = 0.15)& theme(text = element_text(size = 30, face = "bold"), axis.text = element_text(size = 24, colour = "black"))
```

```
## Warning: Some of the plotted features are from meta.data slot.
## • Please check that `na_cutoff` param is being set appropriately for those
##   features.
```

```
## Warning in SetQuantile(cutoff = max.cutoff[index - 3], data.feature): NAs
## introduced by coercion
## Warning in SetQuantile(cutoff = max.cutoff[index - 3], data.feature): NAs
## introduced by coercion
## Warning in SetQuantile(cutoff = max.cutoff[index - 3], data.feature): NAs
## introduced by coercion
## Warning in SetQuantile(cutoff = max.cutoff[index - 3], data.feature): NAs
## introduced by coercion
```

#### Pseudobulk of mesenchymal fibroblasts from integrated stromal
dataset

##### Pseudobulk by sample

```
load(file = "Combined_stromal.Robj")

DefaultAssay(Combined_stromal) <- "RNA"

Combined_stromal <- NormalizeData(Combined_stromal, normalization.method = "RC", scale.factor = 10000)

Idents(Combined_stromal) <- Combined_stromal$AL_clusters
levels(Combined_stromal)
```

```
## [1] "c0 ACKR1+ Endothelial cell"        "c1 Mesenchymal Fibroblast"        
## [3] "c2 Proinflammatory Fibroblast"     "c3 SMC/myofibroblast"             
## [5] "c4 Secretory-papillary Fibroblast" "c5 Endothelial cell"              
## [7] "c6 Lymphatic Endothelial cell"     "c7 Proliferating"
```

```
idents <- ("c1 Mesenchymal Fibroblast")

# Use a loop to subset the data and store in a list
subset_list <- lapply(idents, function(ident) {
  subset(x = Combined_stromal, idents = ident)
})

#name the list for easier access
names(subset_list) <- idents

Idents(Combined_stromal) <- Combined_stromal$orig.ident
levels(Combined_stromal)
```

```
##  [1] "Kd1"      "Kd2"      "Kd3"      "Kd4"      "Nsk1"     "Nsc1"    
##  [7] "Nsc2"     "Nsc3"     "KF1"      "KF2"      "KF3"      "NF1"     
## [13] "NF2"      "NF3"      "K007CASE" "K007CTRL" "K009CASE" "K009CTRL"
## [19] "K012CASE" "K012CTRL" "K013CASE" "K013CTRL"
```

```
idents_second <- c("Kd1", "Kd2", "Kd3", "Kd4", "KF1", "KF2", "KF3", "K007CASE", "K007CTRL", "K009CASE", "K009CTRL", "K012CASE", "K012CTRL", "K013CASE", "K013CTRL")


subset_list_second <- lapply(subset_list, function(first_subset) {
  # Set identities to orig.ident
  Idents(first_subset) <- first_subset$orig.ident
  
  # Find valid identities in the current subset
  valid_idents <- intersect(idents_second, unique(Idents(first_subset)))
  
  # Warn about missing identities
  missing_idents <- setdiff(idents_second, valid_idents)
  if (length(missing_idents) > 0) {
    message("Missing identities in this subset: ", paste(missing_idents, collapse = ", "))
  }
  
  # Initialize the second layer with valid subsets
  second_layer <- lapply(valid_idents, function(ident) {
    subset(x = first_subset, idents = ident)
  })
  
  # Name the second layer by valid sample names
  if (length(valid_idents) > 0) {
    names(second_layer) <- valid_idents
  }
  
  return(second_layer)
})
```

###### Turn each into a matrix and sum the counts for each gene - repeat for all files

```
#define the function
process_seurat <- function(seurat_obj) {
  # Get Assay Data
  data_matrix <- GetAssayData(seurat_obj, slot = "data")
  # Convert to matrix
  data_matrix <- as.matrix(data_matrix)
  # Calculate row sums
  row_sums <- rowSums(data_matrix)
  return(row_sums)
}
# Apply the function to all Seurat objects in the nested list
result_list_nested <- lapply(subset_list_second, function(inner_list) {
  lapply(inner_list, process_seurat)
})
```

```
## Warning in asMethod(object): sparse->dense coercion: allocating vector of size
## 1.6 GiB
```

```
## Warning in asMethod(object): sparse->dense coercion: allocating vector of size
## 1.1 GiB
```

```
# Flatten the nested list into a single-level list with combined names
flattened_result_list <- unlist(result_list_nested, recursive = FALSE)

#combine all the rowSums file into one and export. 
samples_matrix <- do.call(cbind, flattened_result_list)

write.table(samples_matrix, file = "pseudobulk_mesenchymal_by_sample.txt")
#Feed into DESeq2 package for DE analysis
```

#### CellChat

##### Start with P4-only data (all pipeline can be repeated for integrated dataset (Combined\_stromal))

###### Convert to CellChat object

```
# Select the SCT assay
data.input <- GetAssayData(P4, assay = "SCT", slot = "data") 

# Create the CellChat object
cellchat <- createCellChat(object = data.input, meta = P4@meta.data, group.by = "paper_clusters")
```

```
## Warning in createCellChat(object = data.input, meta = P4@meta.data, group.by = "paper_clusters"): The 'meta' data does not have a column named `samples`. We now add this column and all cells are assumed to belong to `sample1`!
```

###### Set human database

```
CellChatDB <- CellChatDB.human 
showDatabaseCategory(CellChatDB)
```

```
CellChatDB.use <- CellChatDB
# set the used database in the object
cellchat@DB <- CellChatDB.use
```

###### Preprocessing the expression data for cell-cell communication analysis

```
cellchat <- subsetData(cellchat)

#Identify over-expressed ligands or receptors in each cell group to infer the cell state-specific communications.
future::plan("multisession", workers = 4)
cellchat <- identifyOverExpressedGenes(cellchat)

#For each overexpressed ligand and receptor obtained above, identify over-expressed L–R interactions if either its associated ligand or receptor is over expressed:
options(future.globals.maxSize = 3 * 1024^3) # Set limit to 2 GiB
cellchat <- identifyOverExpressedInteractions(cellchat)
```

###### Inference of cell–cell communication networks

```
# Remove all objects except  CellChat object
rm(list = setdiff(ls(), "cellchat"))
# Run garbage collection to clear memory
gc()

options(future.globals.maxSize = 8 * 1024^3)
cellchat <- computeCommunProb(cellchat, type = "triMean", trim = NULL, raw.use = TRUE, nboot = 20)
```

```
## Loading required package: dplyr
```

```
## 
## Attaching package: 'dplyr'
```

```
## The following object is masked from 'package:Biobase':
## 
##     combine
```

```
## The following objects are masked from 'package:BiocGenerics':
## 
##     combine, intersect, setdiff, union
```

```
## The following objects are masked from 'package:stats':
## 
##     filter, lag
```

```
## The following objects are masked from 'package:base':
## 
##     intersect, setdiff, setequal, union
```

```
## Loading required package: igraph
```

```
## 
## Attaching package: 'igraph'
```

```
## The following objects are masked from 'package:dplyr':
## 
##     as_data_frame, groups, union
```

```
## The following objects are masked from 'package:future':
## 
##     %->%, %<-%
```

```
## The following objects are masked from 'package:BiocGenerics':
## 
##     normalize, path, union
```

```
## The following objects are masked from 'package:stats':
## 
##     decompose, spectrum
```

```
## The following object is masked from 'package:base':
## 
##     union
```

```
## Loading required package: ggplot2
```

```
## Loading required package: dplyr
```

```
## 
## Attaching package: 'dplyr'
```

```
## The following object is masked from 'package:Biobase':
## 
##     combine
```

```
## The following objects are masked from 'package:BiocGenerics':
## 
##     combine, intersect, setdiff, union
```

```
## The following objects are masked from 'package:stats':
## 
##     filter, lag
```

```
## The following objects are masked from 'package:base':
## 
##     intersect, setdiff, setequal, union
```

```
## Loading required package: igraph
```

```
## 
## Attaching package: 'igraph'
```

```
## The following objects are masked from 'package:dplyr':
## 
##     as_data_frame, groups, union
```

```
## The following objects are masked from 'package:future':
## 
##     %->%, %<-%
```

```
## The following objects are masked from 'package:BiocGenerics':
## 
##     normalize, path, union
```

```
## The following objects are masked from 'package:stats':
## 
##     decompose, spectrum
```

```
## The following object is masked from 'package:base':
## 
##     union
```

```
## Loading required package: ggplot2
```

```
## Loading required package: dplyr
```

```
## 
## Attaching package: 'dplyr'
```

```
## The following object is masked from 'package:Biobase':
## 
##     combine
```

```
## The following objects are masked from 'package:BiocGenerics':
## 
##     combine, intersect, setdiff, union
```

```
## The following objects are masked from 'package:stats':
## 
##     filter, lag
```

```
## The following objects are masked from 'package:base':
## 
##     intersect, setdiff, setequal, union
```

```
## Loading required package: igraph
```

```
## 
## Attaching package: 'igraph'
```

```
## The following objects are masked from 'package:dplyr':
## 
##     as_data_frame, groups, union
```

```
## The following objects are masked from 'package:future':
## 
##     %->%, %<-%
```

```
## The following objects are masked from 'package:BiocGenerics':
## 
##     normalize, path, union
```

```
## The following objects are masked from 'package:stats':
## 
##     decompose, spectrum
```

```
## The following object is masked from 'package:base':
## 
##     union
```

```
## Loading required package: ggplot2
```

```
## Loading required package: dplyr
```

```
## 
## Attaching package: 'dplyr'
```

```
## The following object is masked from 'package:Biobase':
## 
##     combine
```

```
## The following objects are masked from 'package:BiocGenerics':
## 
##     combine, intersect, setdiff, union
```

```
## The following objects are masked from 'package:stats':
## 
##     filter, lag
```

```
## The following objects are masked from 'package:base':
## 
##     intersect, setdiff, setequal, union
```

```
## Loading required package: igraph
```

```
## 
## Attaching package: 'igraph'
```

```
## The following objects are masked from 'package:dplyr':
## 
##     as_data_frame, groups, union
```

```
## The following objects are masked from 'package:future':
## 
##     %->%, %<-%
```

```
## The following objects are masked from 'package:BiocGenerics':
## 
##     normalize, path, union
```

```
## The following objects are masked from 'package:stats':
## 
##     decompose, spectrum
```

```
## The following object is masked from 'package:base':
## 
##     union
```

```
## Loading required package: ggplot2
```

```
#Filter the cell–cell communication, based on the number of cells in each group. By default, the minimum number of cells required in each cell group for cell–cell communication is 10.

cellchat <- filterCommunication(cellchat, min.cells = 10)

cellchat <- computeCommunProbPathway(cellchat)
```

###### Calculate the aggregated cell–cell communication network

```
cellchat <- aggregateNet(cellchat)

#Compute the network centrality scores of the inferred cell–cell communication network.

cellchat <- netAnalysis_computeCentrality(cellchat, slot.name = "netP")
```

```
## Warning: UNRELIABLE VALUE: One of the 'future.apply' iterations
## ('future_sapply-1') unexpectedly generated random numbers without declaring so.
## There is a risk that those random numbers are not statistically sound and the
## overall results might be invalid. To fix this, specify 'future.seed=TRUE'. This
## ensures that proper, parallel-safe random numbers are produced via a parallel
## RNG method. To disable this check, use 'future.seed = NULL', or set option
## 'future.rng.onMisuse' to "ignore".
```

```
## Warning: UNRELIABLE VALUE: One of the 'future.apply' iterations
## ('future_sapply-2') unexpectedly generated random numbers without declaring so.
## There is a risk that those random numbers are not statistically sound and the
## overall results might be invalid. To fix this, specify 'future.seed=TRUE'. This
## ensures that proper, parallel-safe random numbers are produced via a parallel
## RNG method. To disable this check, use 'future.seed = NULL', or set option
## 'future.rng.onMisuse' to "ignore".
```

```
## Warning: UNRELIABLE VALUE: One of the 'future.apply' iterations
## ('future_sapply-3') unexpectedly generated random numbers without declaring so.
## There is a risk that those random numbers are not statistically sound and the
## overall results might be invalid. To fix this, specify 'future.seed=TRUE'. This
## ensures that proper, parallel-safe random numbers are produced via a parallel
## RNG method. To disable this check, use 'future.seed = NULL', or set option
## 'future.rng.onMisuse' to "ignore".
```

```
## Warning: UNRELIABLE VALUE: One of the 'future.apply' iterations
## ('future_sapply-4') unexpectedly generated random numbers without declaring so.
## There is a risk that those random numbers are not statistically sound and the
## overall results might be invalid. To fix this, specify 'future.seed=TRUE'. This
## ensures that proper, parallel-safe random numbers are produced via a parallel
## RNG method. To disable this check, use 'future.seed = NULL', or set option
## 'future.rng.onMisuse' to "ignore".
```

###### make heatmaps

```
plot1 <- netAnalysis_signalingRole_heatmap(cellchat, width = 5, height = 10, pattern = "incoming")
plot2 <- netAnalysis_signalingRole_heatmap(cellchat, width = 5, height = 10, pattern = "outgoing")
plot1+plot2
```

```
## Warning: Heatmap/annotation names are duplicated: Relative strength
```

#### GSVA analysis

##### reading in TPM from CSV

```
tpm <- read.csv(file="/Users/amylock_soper/Documents/Kings_Postdoc/Papers/P0P4/Bulk sequencing/tpm_raw.csv", header=T)
head(tpm)
```

```
##    gene_symbol NDF_C_1 NDF_C_2 NDF_C_3 NDF_C_4 KDF_C_1 KDF_C_2 KDF_C_3 KDF_C_4
## 1       FLT3LG   32.63   29.95   27.85   30.34   31.17   40.53   30.50   36.36
## 2         ERI1    9.51   12.45    9.75   12.59    9.68   10.75   10.99   12.71
## 3       SAP130   29.45   25.82   26.92   24.35   28.87   30.35   26.33   25.13
## 4 LOC100129098    0.00    0.00    0.00    0.00    0.00    0.06    0.00    0.00
## 5        SPIN1   62.93   60.63   58.51   65.04   66.40   67.36   63.62   58.43
## 6         WNT6    0.00    0.00    0.00    0.00    0.00    0.09    0.00    0.00
##   NDF_AA_1 NDF_AA_2 NDF_AA_3 NDF_AA_4 KDF_AA_1 KDF_AA_2 KDF_AA_3 KDF_AA_4
## 1    39.76    32.78    28.09    30.97    36.35    38.11    35.60    31.47
## 2    10.72    10.48    13.78    12.84    11.29    12.59    12.73    13.30
## 3    25.25    27.61    29.44    25.50    26.70    27.83    27.72    28.69
## 4     0.00     0.00     0.00     0.00     0.00     0.00     0.00     0.00
## 5    64.78    62.77    63.76    59.66    63.24    67.65    53.24    67.01
## 6     0.00     0.16     0.00     0.15     0.00     0.00     0.00     0.00
```

```
#Removing gene_symbol column name
names <- rownames(tpm)
rownames(tpm) <- make.names(names, unique=TRUE)
rownames(tpm) <- tpm$gene_symbol
tpm$gene_symbol <- NULL
head(tpm)
```

```
##              NDF_C_1 NDF_C_2 NDF_C_3 NDF_C_4 KDF_C_1 KDF_C_2 KDF_C_3 KDF_C_4
## FLT3LG         32.63   29.95   27.85   30.34   31.17   40.53   30.50   36.36
## ERI1            9.51   12.45    9.75   12.59    9.68   10.75   10.99   12.71
## SAP130         29.45   25.82   26.92   24.35   28.87   30.35   26.33   25.13
## LOC100129098    0.00    0.00    0.00    0.00    0.00    0.06    0.00    0.00
## SPIN1          62.93   60.63   58.51   65.04   66.40   67.36   63.62   58.43
## WNT6            0.00    0.00    0.00    0.00    0.00    0.09    0.00    0.00
##              NDF_AA_1 NDF_AA_2 NDF_AA_3 NDF_AA_4 KDF_AA_1 KDF_AA_2 KDF_AA_3
## FLT3LG          39.76    32.78    28.09    30.97    36.35    38.11    35.60
## ERI1            10.72    10.48    13.78    12.84    11.29    12.59    12.73
## SAP130          25.25    27.61    29.44    25.50    26.70    27.83    27.72
## LOC100129098     0.00     0.00     0.00     0.00     0.00     0.00     0.00
## SPIN1           64.78    62.77    63.76    59.66    63.24    67.65    53.24
## WNT6             0.00     0.16     0.00     0.15     0.00     0.00     0.00
##              KDF_AA_4
## FLT3LG          31.47
## ERI1            13.30
## SAP130          28.69
## LOC100129098     0.00
## SPIN1           67.01
## WNT6             0.00
```

```
# tpm = raw tpm values
```

```
#BiocManager::install("GSVA")

library(GSVA)
```

```
## Warning: package 'GSVA' was built under R version 4.4.3
```

```
library(patchwork)

#log normalising to remove skewing by extreme large or small values
tpm_log <- log2(tpm + 1)
tpm_log_mtx <- as.matrix(tpm_log)
```

```
library(ggpattern)


signature_integ <- (list(mesenchymal = c("ASPN", "COMP", "COL1A1", "COL1A2", "CTHRC1", "COL3A1", "OGN", "POSTN", "COL5A1", "FN1",
"SFRP2", "SPARC", "COL12A1", "COL5A2", "HTRA1", "LUM", "MDK", "BGN", "SFRP4", "PTN",
"PRDX4", "AEBP1", "PCOLCE", "COL11A1", "COL14A1", "DPT", "PPIC", "SPON2", "MMP23B", "MRC2",
"COL6A2", "FBLN1", "RCN3", "COPZ2", "ADAM12", "MFAP4", "FAP", "LGALS1", "CCDC80", "MMP2",
"C1QTNF3", "OLFML3", "CERCAM", "CTSK", "THBS2", "COL6A1", "P4HA2", "THBS4", "CHPF", 
"COL9A3", "CADM1", "SDC1", "SLC5A3", "NRG1", "DKK2", "COL10A1", "NPTX2"),
proinflammatory = c("APOD", "APOE", "CFD", "CCL19", "CXCL12", "PTGDS", "MT2A", "CXCL14",
"APOC1", "MGP", "C1R", "COL4A4", "FGFBP2", "C3", "GPM6B", "P2RY14"), 
SMC_myofib = c("ACTA2", "RGS5", "TAGLN", "ID4", "C11orf96", "TPM2", "SYNPO2", "NOTCH3", "MT1A", "MYL9",
"KCNE4", "COL4A1", "STEAP4", "MYH11", "NDUFA4L2", "COL4A2", "CALD1", "MCAM", "HIGD1B", "TPM1",
"MYLK", "CCDC102B", "PDGFRB", "ADAMTS4", "MT2A", "PDGFA", "S100A4", "MT1M", "FILIP1L", "ADRA2A",
"CRISPLD2", "DES", "ENPEP", "PPP1R14A", "CASQ2", "COX4I2", "AVPR1A", "RRAD", "GJA4", "MT1E",
"CD36", "ACTG2"),
sec_pap = c("PTGDS", "ELN", "CFD", "DCN", "APCDD1", "TWIST2", "SFRP2", "CXCL14",
"CCDC80", "DPT", "LTBP1", "PRG4", "CRYAB", "TTR", "WISP2"),
proliferating = c("HIST1H4C", "CENPF", "TOP2A", "MKI67", "PTTG1", "TYMS", "STMN1", "NUSAP1", "CDK1",
"HIST1H1B", "PCLAF", "CDKN3", "TPX2", "GTSE1", "BIRC5", "UBE2C", "TROAP", "AURKB",
"PIMREG", "CCNB1", "CCNB2", "CDC20", "ASPM", "TK1", "CENPE", "NPTX2", "NUF2",
"CKAP2L", "KRT5")))

# Create parameter object for gsva
gsvaPar <- gsvaParam(tpm_log_mtx, signature_integ)
# Run GSVA
gsva_scores <- gsva(gsvaPar, verbose=TRUE)
```

```
## ℹ GSVA version 2.0.7
```

```
## ! 3 genes with constant values throughout the samples
```

```
## ! Genes with constant values are discarded
```

```
## ℹ Calculating GSVA ranks
```

```
## ℹ kcdf='auto' (default)
```

```
## ℹ GSVA dense (classical) algorithm
```

```
## ℹ Row-wise ECDF estimation with Gaussian kernels
```

```
## ℹ Calculating GSVA column ranks
```

```
## ℹ Calculating GSVA scores
```

```
## ✔ Calculations finished
```

```
# label samples by condition by making metadata column
sample_metadata <- data.frame(
  SampleID = colnames(gsva_scores),
  Condition = c("N_C","N_C","N_C","N_C", "K_C", "K_C", "K_C", "K_C", "N_AA","N_AA","N_AA","N_AA","K_AA", "K_AA", "K_AA", "K_AA" )
)

# Set rownames to match gsva_scores columns
rownames(sample_metadata) <- sample_metadata$SampleID


# Create a data frame for plotting
plot_df <- data.frame(
  Score = gsva_scores["mesenchymal", ],
  Condition = sample_metadata$Condition
)
plot_df$Condition <- factor(plot_df$Condition, levels = c("N_C","K_C", "N_AA","K_AA"))

# ggplot violin + jitter
p1 <- ggplot(plot_df, aes(x = Condition, y = Score, fill = Condition, pattern = Condition)) +
  geom_violin_pattern(
    trim = FALSE,
    alpha = 0.6,
    color = NA,
    pattern_density = 0.2,
    pattern_spacing = 0.03,
    pattern_key_scale_factor = 0.6
  ) +
  geom_jitter(aes(color = Condition), width = 0.15, size = 2.5, alpha = 0.8) +
  scale_fill_manual(values = c("darkgrey","magenta", "darkgrey","magenta")) +  # use white or light fill for pattern visibility
  scale_pattern_manual(values = c("none", "none", "stripe", "stripe")) +  # repeat for each condition level
  scale_color_manual(values = c("black", "black", "black", "black")) +
  labs(
    title = "Enrichment of Mesenchymal Fibroblast cluster",
    y = "GSVA score",
    x = "Condition"
  ) +
  theme_minimal() +
  theme(
    legend.position = "none",
    axis.line = element_line(color = "black"),     # Axis lines
    axis.text = element_text(size = 14),           # Axis text size
    axis.title = element_text(size = 16),          # Axis title size
    plot.title = element_text(size = 18, face = "bold")  # Title text size
  )

print(p1)
```

```
## Registered S3 methods overwritten by 'proxy':
##   method               from    
##   print.registry_field registry
##   print.registry_entry registry
```

```
plot_df <- data.frame(
  Score = gsva_scores["proinflammatory", ],
  Condition = sample_metadata$Condition
)
plot_df$Condition <- factor(plot_df$Condition, levels = c("N_C","K_C", "N_AA","K_AA"))

# ggplot violin + jitter
p2 <- ggplot(plot_df, aes(x = Condition, y = Score, fill = Condition, pattern = Condition)) +
  geom_violin_pattern(
    trim = FALSE,
    alpha = 0.6,
    color = NA,
    pattern_density = 0.2,
    pattern_spacing = 0.03,
    pattern_key_scale_factor = 0.6
  ) +
  geom_jitter(aes(color = Condition), width = 0.15, size = 2.5, alpha = 0.8) +
  scale_fill_manual(values = c("darkgrey","magenta", "darkgrey","magenta")) +  # use white or light fill for pattern visibility
  scale_pattern_manual(values = c("none", "none", "stripe", "stripe")) +  # repeat for each condition level
  scale_color_manual(values = c("black", "black", "black", "black")) +
  labs(
    title = "Enrichment of Proinflammatory Fibroblast cluster",
    y = "GSVA score",
    x = "Condition"
  ) +
  theme_minimal() +
  theme(
    legend.position = "none",
    axis.line = element_line(color = "black"),     # Axis lines
    axis.text = element_text(size = 14),           # Axis text size
    axis.title = element_text(size = 16),          # Axis title size
    plot.title = element_text(size = 18, face = "bold")  # Title text size
  )

print(p2)
```

```
plot_df <- data.frame(
  Score = gsva_scores["SMC_myofib", ],
  Condition = sample_metadata$Condition
)
plot_df$Condition <- factor(plot_df$Condition, levels = c("N_C","K_C", "N_AA","K_AA"))


p3 <- ggplot(plot_df, aes(x = Condition, y = Score, fill = Condition, pattern = Condition)) +
  geom_violin_pattern(
    trim = FALSE,
    alpha = 0.6,
    color = NA,
    pattern_density = 0.2,
    pattern_spacing = 0.03,
    pattern_key_scale_factor = 0.6
  ) +
  geom_jitter(aes(color = Condition), width = 0.15, size = 2.5, alpha = 0.8) +
  scale_fill_manual(values = c("darkgrey","magenta", "darkgrey","magenta")) +  # use white or light fill for pattern visibility
  scale_pattern_manual(values = c("none", "none", "stripe", "stripe")) +  # repeat for each condition level
  scale_color_manual(values = c("black", "black", "black", "black")) +
  labs(
    title = "Enrichment of SMC/Myofibroblast cluster",
    y = "GSVA score",
    x = "Condition"
  ) +
  theme_minimal() +
  theme(
    legend.position = "none",
    axis.line = element_line(color = "black"),     # Axis lines
    axis.text = element_text(size = 14),           # Axis text size
    axis.title = element_text(size = 16),          # Axis title size
    plot.title = element_text(size = 18, face = "bold")  # Title text size
  )
print(p3)
```

```
plot_df <- data.frame(
  Score = gsva_scores["sec_pap", ],
  Condition = sample_metadata$Condition
)
plot_df$Condition <- factor(plot_df$Condition, levels = c("N_C","K_C", "N_AA","K_AA"))


p4 <- ggplot(plot_df, aes(x = Condition, y = Score, fill = Condition, pattern = Condition)) +
  geom_violin_pattern(
    trim = FALSE,
    alpha = 0.6,
    color = NA,
    pattern_density = 0.2,
    pattern_spacing = 0.03,
    pattern_key_scale_factor = 0.6
  ) +
  geom_jitter(aes(color = Condition), width = 0.15, size = 2.5, alpha = 0.8) +
  scale_fill_manual(values = c("darkgrey","magenta", "darkgrey","magenta")) +  # use white or light fill for pattern visibility
  scale_pattern_manual(values = c("none", "none", "stripe", "stripe")) +  # repeat for each condition level
  scale_color_manual(values = c("black", "black", "black", "black")) +
  labs(
    title = "Enrichment of Sec.Papillary Fibroblast cluster",
    y = "GSVA score",
    x = "Condition"
  ) +
  theme_minimal() +
  theme(
    legend.position = "none",
    axis.line = element_line(color = "black"),     # Axis lines
    axis.text = element_text(size = 14),           # Axis text size
    axis.title = element_text(size = 16),          # Axis title size
    plot.title = element_text(size = 18, face = "bold")  # Title text size
  )

print(p4)
```

```
plot_df <- data.frame(
  Score = gsva_scores["proliferating", ],
  Condition = sample_metadata$Condition
)
plot_df$Condition <- factor(plot_df$Condition, levels = c("N_C","K_C", "N_AA","K_AA"))


p5 <- ggplot(plot_df, aes(x = Condition, y = Score, fill = Condition, pattern = Condition)) +
  geom_violin_pattern(
    trim = FALSE,
    alpha = 0.6,
    color = NA,
    pattern_density = 0.2,
    pattern_spacing = 0.03,
    pattern_key_scale_factor = 0.6
  ) +
  geom_jitter(aes(color = Condition), width = 0.15, size = 2.5, alpha = 0.8) +
  scale_fill_manual(values = c("darkgrey","magenta", "darkgrey","magenta")) +  # use white or light fill for pattern visibility
  scale_pattern_manual(values = c("none", "none", "stripe", "stripe")) +  # repeat for each condition level
  scale_color_manual(values = c("black", "black", "black", "black")) +
  labs(
    title = "Enrichment of Proliferating Fibroblast cluster",
    y = "GSVA score",
    x = "Condition"
  ) +
  theme_minimal() +
  theme(
    legend.position = "none",
    axis.line = element_line(color = "black"),     # Axis lines
    axis.text = element_text(size = 14),           # Axis text size
    axis.title = element_text(size = 16),          # Axis title size
    plot.title = element_text(size = 18, face = "bold")  # Title text size
  )
print(p5)
```

```
combined_plots <- (p1 | p2 | p3 | p4 | p5) + plot_layout(ncol = 3, nrow = 2)
print(combined_plots)
```
